# Absolute Scaling of Single-Cell Transcriptomes Reveals Pervasive Hypertranscription in Adult Stem and Progenitor Cells

**DOI:** 10.1101/2021.12.13.472426

**Authors:** Yun-Kyo Kim, Miguel Ramalho-Santos

## Abstract

Hypertranscription facilitates biosynthetically demanding cellular state transitions through global upregulation of the nascent transcriptome. Despite its potential widespread relevance, documented examples of hypertranscription remain few and limited predominantly to early development. This limitation is in large part due to the fact that modern sequencing approaches, including single-cell RNA sequencing (scRNA-seq), generally assume similar levels of transcriptional output per cell. Here, we use molecule counting and spike-in normalization to develop absolute scaling of single-cell RNA sequencing data. Absolute scaling enables an estimation of total transcript abundances per cell, which we validate in embryonic stem cell (ESC) and germline data and apply to adult mouse organs at steady-state or during regeneration. The results reveal a remarkable dynamic range in transcriptional output among adult cell types. We find that many different multipotent stem and progenitor cell populations are in a state of hypertranscription, including in the hematopoietic system, intestine and skin. Hypertranscription marks cells with multilineage potential in adult organs, is redeployed in conditions of tissue injury, and can precede by 1-2 days bursts of proliferation during regeneration. In addition to the association between hypertranscription and the stem/progenitor cell state, we dissect the relationship between transcriptional output and cell cycle, ploidy and secretory behavior. Our analyses reveal a common set of molecular pathways associated with hypertranscription across adult organs, including chromatin remodeling, DNA repair, ribosome biogenesis and translation. Our findings introduce an approach towards maximizing single-cell RNA-seq profiling. By applying this methodology across a diverse collection of cell states and contexts, we put forth hypertranscription as a general and dynamic cellular program that is pervasively employed during development, organ maintenance and regeneration.

**SUMMARY STATEMENT:** Absolute scaling of single-cell transcriptomic data reveals highly dynamic global levels of transcription across adult cell lineages

## INTRODUCTION

Cells dynamically regulate their biosynthetic capacities to fulfill the requirements of demanding cell state transitions^1^. At the level of transcription, cells can meet these demands by entering a state of relative hypertranscription, which is characterized by a global upregulation of nascent transcriptional output^2^. This global shift powers biological phenomena requiring substantial increases in total biomass, such as rapid proliferation, secretion, and cell activation^1,2^. Hence, hypertranscription has been proposed to play major roles in developmental transitions, adult organ homeostasis, and tumorigenesis^2^. Our group and others have reported evidence supporting the occurrence and functional relevance of hypertranscription in mouse embryogenesis and select human cancer cell types^3–8^. However, the occurrence of hypertranscription in adult physiology remains largely unexplored

Despite the paucity of data in adult tissues, work in mouse embryos and mESCs has identified five recurring “hallmarks” of hypertranscription. First, hypertranscribing cells display a transcriptome-wide increase in gene expression, including at housekeeping genes and ribosomal RNA (rRNA)^4,9^. These changes occur at the level of both steady-state and nascent transcripts, and result in measurable increases in total cellular RNA content^4^. Second, hypertranscription is dependent on a decondensed and distinctly permissive chromatin landscape, maintained by the activity of euchromatic remodeling factors^10,11^. One notable factor is the ATP-dependent remodeler Chd1, which binds specifically to H3K4me3 and is required for the transcriptional output of proliferating epiblast cells^4^. Third, hypertranscribing cells upregulate protein synthesis/translational machinery, which is necessary for the continuous translation and steady-state maintenance of Chd1 and several other unstable euchromatic regulators^10^. Fourth, hypertranscribing cells endogenously accumulate promoter-proximal double strand breaks (DSBs), thus displaying a heightened dependency on DNA repair factors^12^. Fifth, hypertranscription is thought to be mediated by the expression of general and “universally amplifying” transactivators, of which the Myc family of transcription factors is the best characterized^3,13^.

A critical requirement for detecting hypertranscription is the ability to distinguish absolute differences in transcript expression between samples or cell states. Despite substantial modern advancements in bulk and single-cell transcriptomic profiling, such measurement is complicated by the standard use of between-transcriptome normalization procedures^14^. For instance, normalization to read-depth in RNA-seq data or housekeeping genes in qRT-PCR assume similar amounts of cellular RNA content between samples of interest. Analysis of scRNA-seq commonly employs similar global scaling approaches, which are used to minimize the impact of stochastic technical effects including capture inefficiencies, amplification biases, and variable sequencing depths^14–16^. However, the global scaling tools used to eliminate these technical biases also suppress genuine biological differences in mRNA content, thereby masking the detection of global shifts driven by hypertranscriptive states^14^. Thus, the analysis of hypertranscription at single-cell resolution requires development of alternative methodologies.

In previous work investigating hypertranscription, we circumvented the problem of between-sample normalization by developing cell-number-normalized (CNN) profiling methods which accurately reproduce absolute cellular transcript levels^3,4,9^. Here, we apply an analogous approach to single-cell transcriptomic data by leveraging unique molecular identifiers (UMIs) or External RNA Controls Consortium (ERCCs) spike-in sequences, two tools commonly included in various scRNA-seq protocols^17–19^. We use these tools to perform absolute scaling of scRNA-seq data, which we validate as being able to faithfully capture hypertranscription in ground-truth settings and developmental cases of hypertranscription previously demonstrated using bulk CNN RNA-seq. We apply this methodology to multiple scRNA-seq datasets in heterogenous adult tissues and report the identification of progenitor cell lineages displaying hallmarks of hypertranscription during homeostasis and regeneration. These results support a model where hypertranscription acts as a general mechanism to facilitate dynamic regulation of cellular biosynthetic capacity during development, organ homeostasis, and regeneration.

## RESULTS

### Absolute Scaling Accurately Estimates Transcript Content in Ground Truth Data

To estimate absolute cellular transcript abundances in scRNA-seq data, we focused on protocols using UMIs or ERCCs for normalization and quality control **(Supp Fig 1A)**. ERCC spike-in sequences are typically used in flow cytometric methods (e.g., Smart-seq2, CEL-seq) and are added to lysis buffer solutions at known concentrations. Similar to our previous bulk CNN RNA-seq methodology, the equal distribution of spike-ins prior to capture allows scaling factors to be generated using the ERCC fraction of reads^3^. These scaling factors normalize for downstream technical effects while also retaining biological differences in total transcript abundance^14^. In contrast, UMIs are typically used in droplet and split-pooling methods (10X Genomics, sci-RNA-seq3) and uniquely label each transcript molecule during mRNA capture. Although raw UMI counts retain variation due to per-cell capture efficiency and dropout, we reasoned that they can still provide an acceptable estimate of transcript abundance when analyzed across large enough samples of cells. Collectively, we refer to these approaches to normalization as *absolute scaling*, in contrast to the *global scaling* performed in typical normalization of scRNA-seq data **(Supp Fig 1B)**.

To evaluate whether UMIs and ERCCs can be used for absolute scaling, we explored two ground truth scenarios. Firstly, we assessed whether using raw UMI counts can distinguish differences in total transcript abundance from UMI-containing libraries generated from singlets vs doublets of cells. As doublet libraries are expected to contain roughly twice the mRNA of singlets, they represent one scenario of hypertranscription with mostly uniform amplification across the transcriptome. We used three publicly available 10X Genomics datasets with experimental doublet annotation: in two datasets based on peripheral blood mononuclear cells (PBMCs), we found that doublets contained roughly 1.8 to 1.9 times the total transcripts as singlets, consistent with previous reports **(Supp Fig 2A)**^20,21^. These doublets also displayed similar increases in the expression of ribosomal and housekeeping genes **(Supp Fig 2B)**. In mixtures of different cell types, such as human and mouse, transcript content of singlets was reflective of known cell size differences (17946.0 UMIs in 293T vs 10522.0 UMIs in 3T3)^22^. Importantly, transcripts in doublets were consistent with aggregation of these two cell types (mean of 26222.5 UMIs), indicating accurate capture of total abundance **(Supp Fig 2A)**.

Secondly, we assessed whether raw UMIs and ERCCs can recapitulate mRNA differences in artificially generated pseudo-cell libraries. We took advantage of the previously published “scRNA-seq Mixology” experiment, which generated UMI and ERCC-containing CEL/Sort-seq libraries using extracted mRNA over a 4-step dilution series^23^. This series contained material from three different human cell lines and spanned an order of magnitude, sufficiently representing the variation found in hypertranscribing cells^3,4^. As the mRNA content of each single-cell library was known, we evaluated the reproduction of transcript abundance using either raw total UMIs or ERCC-derived per-cell size factors. Compared to global scaling, absolute scaling using either method more accurately reproduced ground truth differences in pseudo-cell transcript levels. **(Supp Fig 3A, C)**. Importantly, changes in transcript abundances were seen to be largely uniform across detected genes, analogous to what would be observed in biological hypertranscription **(Supp Fig 3B, D)**. We found that regardless of platform, ERCC normalization slightly outperformed raw UMI counts and generated lower inter-cell variability in transcript abundances, particularly at higher ground truth RNA amounts **(Supp Fig 3A-D, E-F)**. Together, these results document the utility of ERCCs and UMIs in absolute scaling and recovering global transcriptomic differences in scRNA-seq data.

### Validation of Absolute Scaling Using Embryonic Hypertranscription Data

We next sought to test absolute scaling in previously established contexts of hypertranscription. The early mouse embryo is especially well characterized in this regard and can be modeled in-vitro using mESCs^24^. Specifically, serum/LIF-grown mESCs are hypertranscriptional and represent the rapidly proliferating early post-implantation epiblast, while mESCs under dual GSK/MEK inhibition (2i media) are comparatively hypotranscriptional and represent a pre-implantation-like state^9,24^. We performed absolute scaling on an ERCC-spiked mESC dataset generated using Fluidigm C1 containing both serum and 2i cells^25^. While globally-scaled data showed only a modest 1.02-fold change between the two cell states, ERCC-normalization revealed that serum mESCs contain 2.37-fold higher transcript counts than 2i mESCs, in concordance with findings from bulk CNN RNA-seq data **(Fig 1A-F)**^9^.

**Figure 1:**
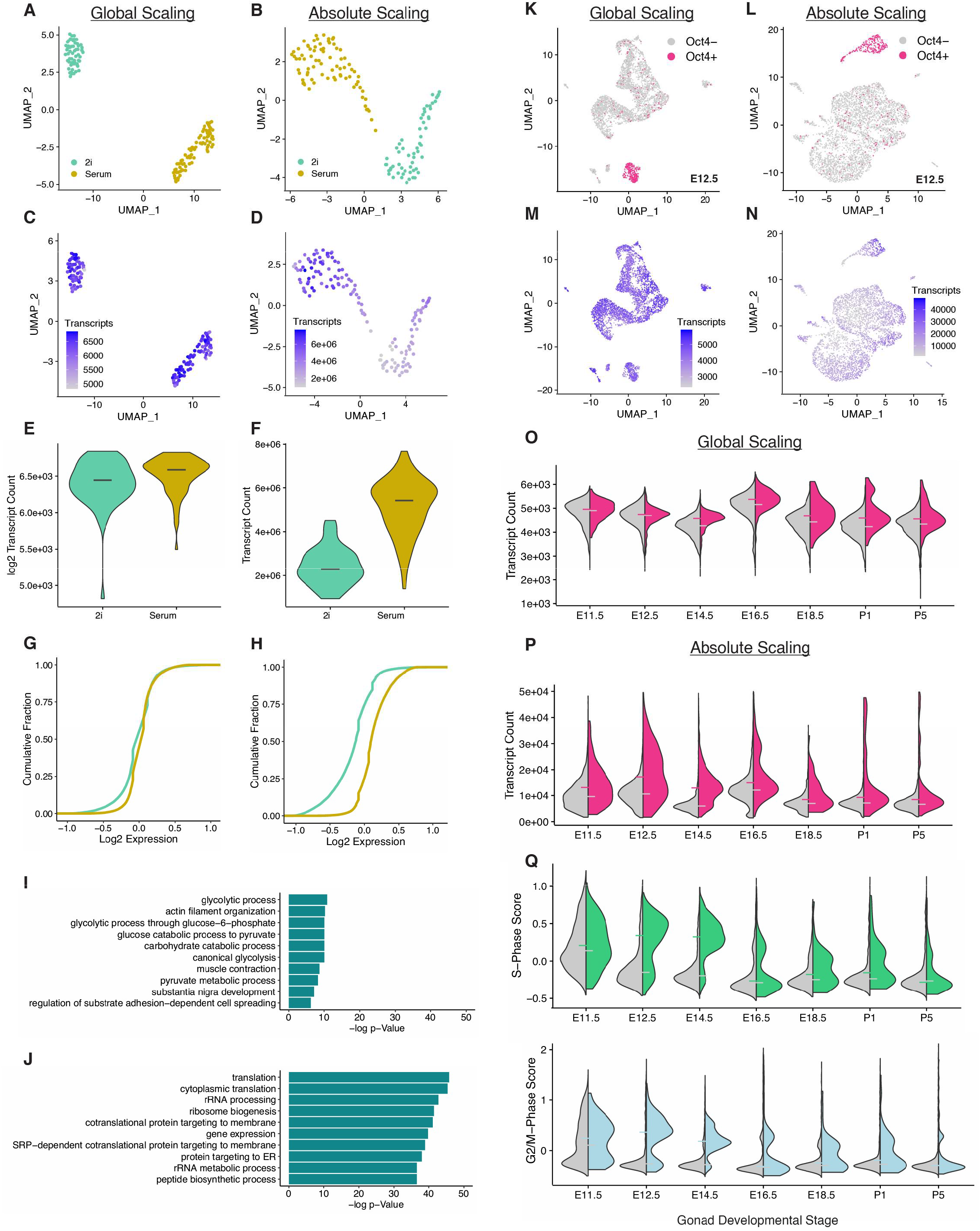
Detection of Embryonic Hypertranscription by Absolute Scaling. **(A-B)** UMAP visualization of serum/2i mESCs under global/absolute scaling. Plots depict dimensionality reduction performed under the indicated scaling type. **(C-F)** Total transcript counts in serum/2i mESCs under global/absolute scaling. Counts in **(C)**, **(E)** represent sums of globally scaled transcripts following log2 scaling, while counts in **(D)**, **(F)** represent total transcripts following ERCC normalization. Lines correspond to median values. **(G-H)** Cumulative distribution plots depicting gene expression across top 25,000 genes in serum/2i mESCs. Distributions were computed using log2 expression in both conditions. **(I-J)** Enriched GO terms from ≥2-fold differentially upregulated genes in serum mESCs under global or absolute scaling conditions. **(K-L)** UMAP visualization of E12.5 gonad scRNA-seq data under global/absolute scaling containing Oct4+ PGCs and Oct4-soma. Plots depict dimensionality reduction performed under the indicated scaling type. **(M-P)** Transcript abundance in gonad scRNA-seq data under global/absolute scaling. Counts in **(M)**, **(O)** represent sums of globally scaled UMIs following log2 scaling, while counts in **(N)**, **(P)** represent raw UMIs. Lines correspond to median values and are coloured according to cell type. **(Q)** Cell cycle scoring of gonad cells. Phase scores were calculated using cell cycle markers from globally-scaled gene expression data. Lines correspond to median values and are coloured according to cell type.

To confirm whether this increase in total transcript abundance is reflective of hypertranscription, we turned to evaluating known hallmarks. We found that under absolute scaling, mESCs display global transcriptomic upregulation across both highly and lowly expressed genes **(Fig 1G-H)**. These upregulated genes include chromatin remodelers, DNA repair factors, ribosomal genes, housekeeping genes, and *Myc* **(Supp Fig 4 A-B)**. Performing Gene Ontology (GO) analysis using differentially expressed genes in serum vs 2i mESCs extracted from globally scaled data revealed an enrichment for terms associated predominantly with metabolic pathways, mainly highlighting the metabolic differences between the cell states. **(Fig 1I)**. In contrast, GO analysis starting with absolute scaling revealed a high enrichment of terms related to translational/ribosomal processes and biosynthesis **(Fig 1J)**. These findings are in strong agreement with our previous bulk-CNN characterizations of hypertranscribing serum vs 2i mESCs^9^.

We then looked to validate our absolute scaling methodology using data from cells in vivo, rather than cultured cells. We have previously shown that during mouse embryogenesis, the mid-gestation primordial germ cell (PGC) lineage undergoes Myc-driven hypertranscription relative to the surrounding soma^3^. To assess whether hypertranscription can also be captured during this period in scRNA-seq data, we used a 10X Genomics dataset generated from whole mouse gonads at seven timepoints between E11.5 to P5^26^. When identified by expression of *Oct4*, globally scaled PGCs appear as having minimal difference in total transcript counts compared to soma **(Fig1 K, M, O)**. In contrast, an analysis of raw UMIs revealed that PGC transcript abundances are 1.62-fold and 2.17-fold increased over soma at E12.5 and E14.5, respectively **(Fig 1L, N, P)**. These elevations in UMI counts correspond precisely to the timepoints of PGC mitotic expansion during development and are accompanied by increases in G2/M and S-phase cell cycle scores **(Fig 1Q)**^26,27^.

Similar to serum mESCs, the transcriptomes of E12.5 and E14.5 PGCs normalized by absolute scaling display a high overrepresentation of genes related to hypertranscription hallmarks **(Supp Fig 4C-D)**. Importantly, these differential expression changes are largely absent prior to and following the burst in PGC proliferation, in agreement with the restriction of hypertranscription to mid-gestation development^3^. Together, these data indicate that absolute scaling, whether based on UMI or ERCC-spiked data, accurately reproduces embryonic hypertranscription in scRNA-seq data both in culture and in vivo, prompting us to apply it to complex adult cell datasets.

### Transcriptional Content Heterogeneity Across Adult Cell Lineages

We previously hypothesized that the rapid proliferation required for turnover in select organs systems may be facilitated in part by hypertranscription^2^. We looked to explore this possibility by applying the absolute scaling methodology described above to existing single-cell atlases of mouse organs. Large-scale scRNA-seq studies have begun to assemble compendia of cellular diversity, which serve as general resources of organism-wide cell expression profiles^28–30^. We chose to focus on the Tabula Muris atlas of twenty mouse organs, which was particularly amenable to absolute scaling due to its inclusion of both ERCC spike-ins and UMIs^31^. Within the atlas, these inclusions are divided between datasets generated using Smart-seq2 (FACS) and 10X Genomics (Droplet) platforms, respectively. In the following analysis, we focused primarily on FACS datasets due to the wider organ selection and greater read-depth, as well as the better performance of ERCC normalization for absolute scaling (see **Supp Fig 3**)^14^. Where relevant, we corroborated significant findings using data based on raw UMI counts.

To apply absolute scaling to the Tabula Muris, we first ensured linear correlations of ERCC expression between organ datasets, as well as validated that individual ERCC species were present at expected frequencies **(Supp Fig 5G,I)**. In addition, we performed filtering for ERCC levels, library size, and doublets for all datasets **(Supp Fig 5H**, and see Methods)^32^. These steps resulted in a total of 42,047 ERCC-containing and 45568 UMI-containing libraries, which upon absolute scaling showed an interquartile range (IQR) of 1,119,112 transcripts and 6377 UMIs, respectively **(Supp Fig 5A-D)**. Where available, FACS and Droplet datasets of the same organ were correlated at the level of both gene expression and median transcripts per cell **(Supp Fig 5J-K)**.

Given the substantial IQR of cellular transcript content in both FACS and Droplet datasets, we next looked to assess whether transcript content differed significantly at the level of individual organ systems. Remarkably, we found that the Tabula Muris displays substantial variation in median transcript counts per organ, spanning a 6.63-fold difference in FACS data and a 3.78-fold difference in 10X data **(Fig 2A, Supp Fig 6A, Table S1)**. The distribution of transcript abundances within organs also varies substantially, with the highest variance in the large intestine (FACS IQR = 2,454,958) or bladder (10X IQR = 10987) and lowest variance in the spleen (Smartseq 2 IQR =199384, 10X IQR = 2174) **(Fig 2A, Supp Fig 6A-B, Table S1)**. Interestingly, we noticed that datasets derived from the hematopoietic, intestinal, and integumentary organs – systems classically associated with continuous turnover^33^ – rank highly amongst other organs in transcript content and are bimodally distributed (see below).

**Figure 2:**
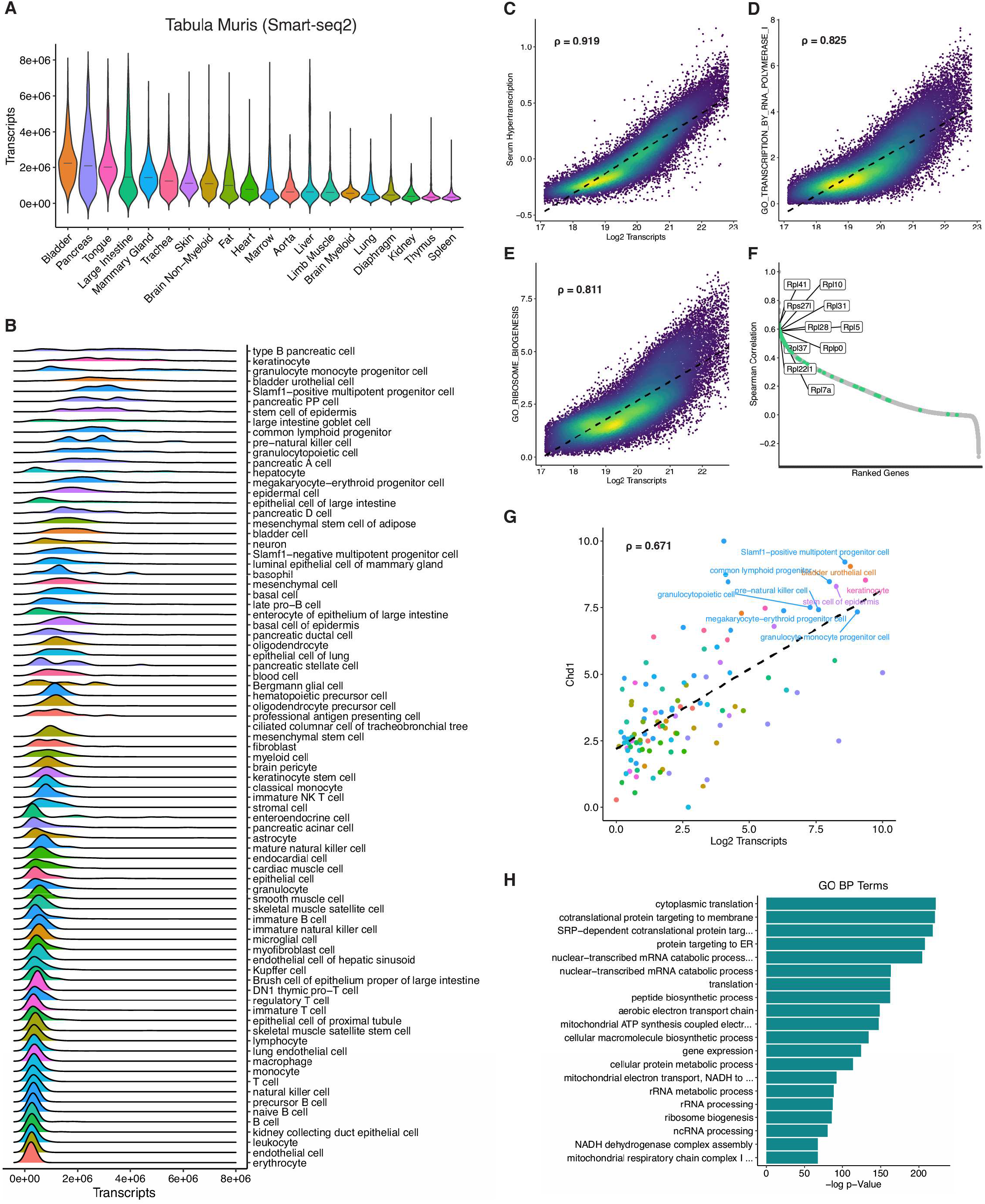
Absolute Scaling Reveals Transcriptional Content Heterogeneity in the Tabula Muris. **(A-B)** Distribution of cellular transcript content between FACS organ datasets under absolute scaling, ranked by median total transcripts. Transcripts represent ERCC-normalized reads, lines correspond to median values. **(C-E)** Single-cell correlation of transcription signatures with log2 transcript abundance. Signature scores were determined using VISION with absolute-scaled FACS data across the entire atlas. Each point represents a single cell, with color scale depicting plotting density. **(F)** Ranking of genes by Spearman coefficient to log2 transcript abundance across the entire FACS atlas. Highlighted genes represent all expressed Rpl and Rps genes. **(G)** Correlation between *Chd1* expression with log2 transcript abundance. Each point is representative of a single averaged cell type, with colors matching organs in **(A)**. Correlation value represents Spearman coefficient. **(H)** Enrichment analysis for GO biological process terms using top 3000 highly correlated genes (to log2 transcript abundance) under absolute scaling.

We next looked to visualize the distribution and heterogeneity of transcript abundance in the Tabula Muris. As absolute scaling would be expected to change the structure of our data (**Fig 1A** vs **B**), we performed dimensionality reduction under global scaling, but visualized transcript counts generated under absolute scaling. We found that even under these conditions of RNA content parity, the distribution of transcript counts are heterogenous and a defining characteristic of many cell clusters **(Supp Fig 7A-B)**. In addition, we observed widespread presence of transcript abundance gradients between highly separated groups of cells, suggesting intermediate states of cellular transcript content. To further explore these differences, we leveraged cell type annotations by the Tabula Muris Consortium, which revealed that transcript counts between different cell types span a remarkable 15.79-fold difference in FACS data (over 82 cell types) and a 8.29-fold difference in 10X data (over 56 cell types) **(Fig 2B, Supp Fig 6B, Table S1)**^31^. Interestingly, we found that a large proportion of the highest transcript-count cells are associated with an adult stem-cell or progenitor-cell identity, particularly from the bone marrow, intestine, and skin datasets. These findings are supported by the observation of higher levels of uridine incorporation in intestinal crypts, where stem and progenitor cells reside (ref^34^ and see below). In contrast, we found that cell-types with the lowest transcript counts are often associated with more highly differentiated or terminal cell states **(Fig 2B, Supp Fig 6B, Table S1)**. Some notable exceptions to this association between elevated transcript counts and stem/progenitor cell states are discussed below. The accuracy of these differences was supported by transcript abundances in T-cells, B-cells, macrophages, and endothelial cells, which together display minimal variation while spanning multiple organ datasets **(Supp Fig 8A-B)**.

Together, these data reveal a remarkable degree of inter- and intra-organ variation with regards to transcript content per cell. As a corollary, these findings suggest that application of global-scaling approaches in complex tissues, while useful for identifying outlier modules of differential expression, are insufficient to capture important biological information with regards to transcriptional output per cell.

### Relationship Between Elevated Transcriptional Output and the Cell Cycle

We next looked to evaluate whether high-content cell types were possible candidates for novel hypertranscribing populations. As hypertranscription is thought to facilitate rapid proliferation in embryonic contexts, we assessed the relationship between cell cycling and transcript abundance^2^. Using globally-scaled expression data, we performed cell cycle staging as previously described^35^ and evaluated the proportion of cycling and non-cycling cells within each annotated cell identity. Surprisingly, when identities were ranked by the proportion of non-cycling cells, we observed a roughly bimodal distribution with regards to total transcript content **(Supp Fig 9, Supp Fig 10)**. Cell types canonically associated with rapid turnover, including progenitor cells of the bone marrow, skin, and intestines, display high proportions of G2/M and S-phase cells alongside elevated transcript counts. In contrast, some non-cycling G1 cells of the liver and bladder urothelium also display heightened transcript abundance.

We hypothesized that the properties of these latter cell types might in part be driven by alternative mechanisms, including somatic polyploidy. Polyploidy has been well documented in both hepatocytes and superficial urothelial cells, and arises through either incomplete cytokinesis or endoreplication^36–41^. Although we lacked DNA content information to directly measure these processes, we took advantage of previous studies identifying markers specific for polyploid cell populations. Using the Tabula Muris FACS liver dataset, we found that cell clusters with elevated transcript abundances are also enriched in markers of 4n hepatocyte *Mixipl*, *Lifr*, and *Nr1i3*^42^ **(Supp Fig 11A-D)**. Similarly, we found that high-transcript cells within the FACS bladder dataset co-express *Krt20* and *Upk2* while lacking *Trp63*, a combination specific for 4n+4n superficial cells^36^ **(Supp Fig 11E-O)**. Thus, these data suggest that estimation of mRNA abundance using absolute scaling is consistent with expected differences driven by polyploidy^38^.

A further exception to the general association between high transcript content and cycling stem/progenitor cells appears to be secretory terminally differentiated cells, such as goblet and Paneth cells within the intestinal epithelium **(Supp Fig 14**, see below). We had previously speculated that the biosynthetic demand of secretory cells may require features of hypertranscription^2^. This possibility is supported by the present analysis but warrants further investigation. Taken together, these analyses indicates that elevated transcriptional counts are generally associated with cycling stem/progenitor cells across multiple organs, with important exceptions observed in terminally differentiated cells that may be in part explained by polyploidy or high secretory output.

### Adult Cells with High Transcript Content Display Hallmarks of Hypertranscription

We next investigated the relationship between transcript content and hypertranscription using properties previously defined in bulk RNA-seq studies of embryonic cell populations (see Introduction and ref^2^). First, we chose a global approach by evaluating the atlas-wide expression of various transcriptional signatures representative of relevant biological processes. At the level of single cells, we found that transcript abundance is robustly correlated to GO hallmark gene sets previously associated with hypertranscription, such as chromatin organization, RNA Pol II activity, ribosome biogenesis, DNA repair, and cell division **(Fig 2C-E, Supp Fig 12A-C)**. We also found that a signature of embryonic hypertranscription, generated using the earlier Kolodziejczyk et al^25^ dataset, is highly correlated with transcript content in adult cells **(Fig 2C)**. These associations are continuous, suggesting that cells do not endogenously rest at discrete levels of transcription. Rather, these data suggest that transcriptional output is fine-tuned across a wide spectrum by euchromatin regulation, RNA stability, translation, and other related processes.

We next sought to assess whether these hallmarks are reflected at the level of the expression of key individual regulators of hypertranscription. We found that the absolute-scaled expression of *Chd1*, transcription factors of the *Myc* family (see below), several DSB repair factors, mTOR, and ribosomal proteins are robustly correlated with single-cell transcript content **(Fig 2F-G, Supp Fig 12D-I, Table S2)**. Additionally, performing enrichment analysis on the set of genes with correlation coefficients of >=0.75 to transcriptional content per cell revealed strong enrichments for GO terms related to translation, transcription, and metabolic processes **(Fig 2H, Table S3)**.

A key hypothesis underlying the mechanistic basis of hypertranscription is the activity of universally amplifying transcription factors, which bind and regulate large portions of the transcriptome. Experimental evidence for both Myc and Yap-driven hypertranscription have been reported in embryonic contexts^3,7,8^. To explore whether the role of these transcription factors is retained in adult cells, we first used ChIP Enrichment Analysis (ChEA) on the Tabula Muris using genes with a >=0.75 Spearman’s coefficient to transcript abundance. ChEA performs gene-set enrichment using protein-DNA interactions derived from ChIP-seq and DamID studies^43^. Interestingly, we found that genes with strong correlations to transcript content are highly enriched in Myc and N-Myc ChEA interactions across several biological contexts **(Supp Fig 12J, Table S3)**. In contrast, correlated genes display weaker enrichment in Yap interactions **(Table S3)**. These differences are reflected in the overall higher correlation of per-cell transcript content with *Myc* over *Yap* **(Supp Fig 12G-I)**. Thus, the Myc family of transcription factors, more so than Yap, are strongly associated with hypertranscription in adult stem/progenitor cells. This analysis further reveals other transcription factors that have not to date been studied in the context of hypertranscription and display chromatin binding patterns strongly linked to transcript content in adult stem/progenitor cells **(Supp Fig 12J, Table S3)**. These transcription factors, such as Eklf, E2f1, and Zfx, deserve to be revisited for potential roles in hypertranscription.

Taken together, these data demonstrate a strong congruity of hypertranscription hallmarks between established embryonic models and stem/progenitor cells of adult tissues^3–5,7–10,25^. These similarities indicate that adult cycling stem/progenitor cells redeploy an embryonic hypertranscriptional program to meet the biosynthetic demands of organ renewal.

### Hypertranscription Marks Cells with Multilineage Potential in Adult Organs

The presence of hypertranscription hallmarks within adult multipotent stem/progenitor cells led us next to probe the association between transcript content and lineage progression. We first asked whether cellular transcript counts are correlated with any previously established methods of inferring differentiation trajectories in scRNA-seq data. Differentiation progression has been captured using feature counts (number of expressed genes) or quantile polarization scores of principal components of the data (possibly reflecting polarized biological activity), providing the framework for the CytoTRACE and VECTOR trajectory inference tools, respectively^44,45^. Across individual cells of each organ dataset, we found that transcript abundance is well correlated with both metrics, displaying particularly strong associations with feature counts **(Supp Fig 13A-B)**. These data suggest that transcriptional output, alongside transcript diversity, decreases along differentiation trajectories concomitant with a loss of multipotent capacity.

To explore this concept in the context of a well-defined differentiation model, we focused specifically on hematopoiesis within the bone marrow, the progression of which has been extensively characterized^46^. We found that ranking cell-types by absolute transcript abundance approximates the known hematopoietic hierarchy from multipotent stem cells to terminal differentiated mononuclear cell types **(Fig 3A-C)**. Importantly, hematopoietic stem/progenitor cells display the previously explored hallmarks of hypertranscription, including upregulation of chromatin remodelers, DNA repair factors, ribosomal genes, and housekeeping genes **(Fig 3D),** as well as a substantial enrichment of the embryonic stem cell serum hypertranscription signature **(Fig 3E).** Moreover, we found that hematopoietic stem/progenitor cells display higher levels of both steady-state and nascent (intronic) reads, in agreement with the notion that hypertranscription is primarily regulated at the level of nascent transcription^2^ **(Fig 3F)**.

**Figure 3:**
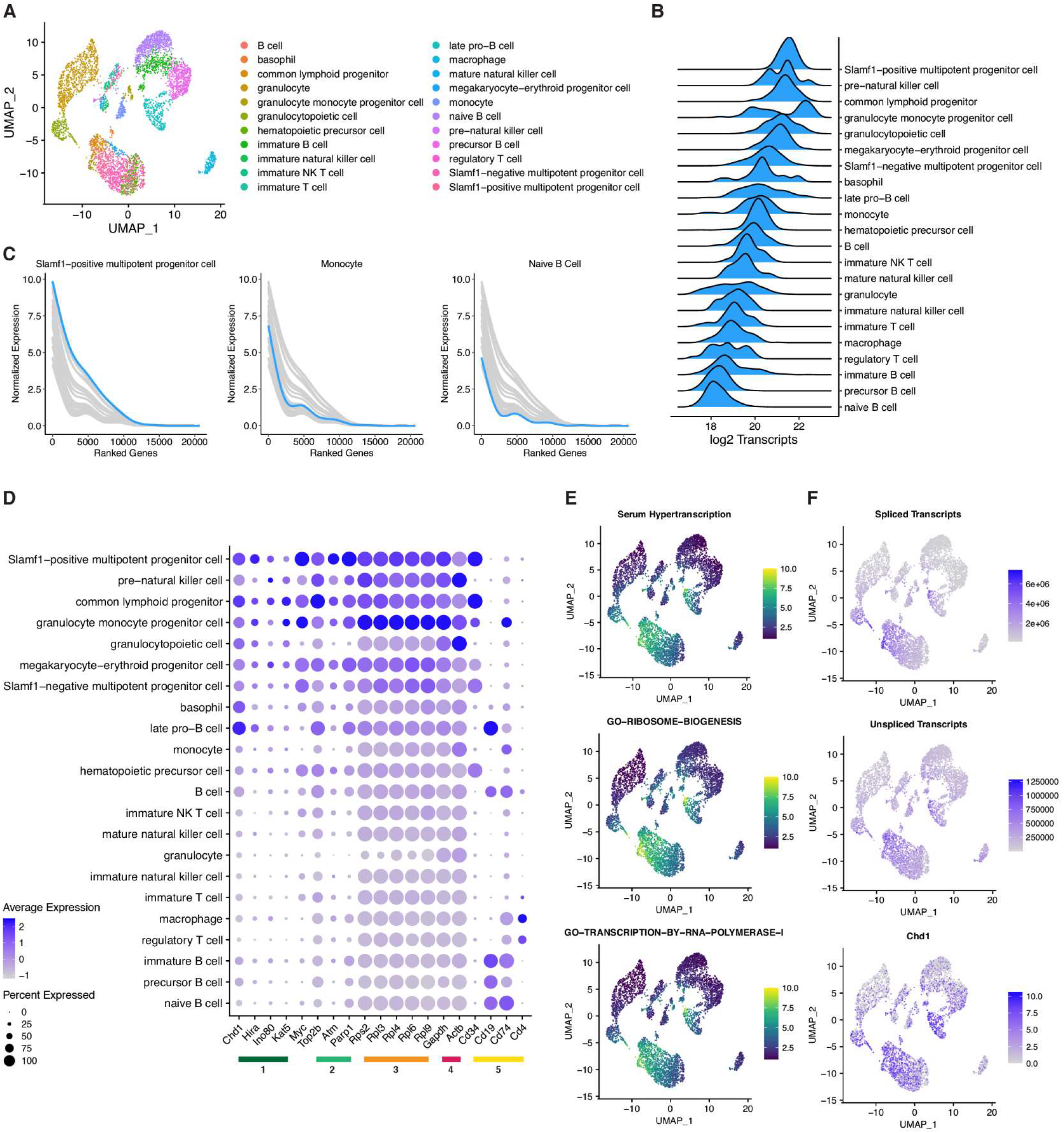
Hematopoietic Progenitors Display Hallmarks of Hypertranscription. **(A)** UMAP visualization of cell types within the bone marrow dataset using dimensionality reduction performed under absolute scaling. **(B)** Distribution of cellular transcript content between cell types of the bone marrow FACS dataset. Transcript counts represent log2 ERCC-normalized reads. **(C)** Transcriptome curves depicting gene expression across top 20,000 genes in representative cell types of the bone marrow. Individual genes are ranked using combined log2 expression between all cell types. Indicated cell types correspond to the highlighted curve. **(D)** Expression of select genes relevant to hallmarks of hypertranscription, including (1) chromatin remodelers, (2) DNA repair factors, (3) ribosomal genes, (4) housekeeping genes, and (5) hematopoietic markers. **(E)** UMAP visualization of signature scores generated by VISION using absolute-scaled expression data. **(F)** UMAP visualization of total spliced/unspliced transcripts and *Chd1* expression under absolute scaling. Unspliced or nascent transcripts were assigned by the detection of intronic sequences.

Consistent with the model that hypertranscription acts in adult progenitors to facilitate renewal, we identified concordant results in stem cells of the skin and adult large intestine. In the interfollicular epidermis, *Top2a*+ cycling basal stem cells contain elevated transcript abundance compared to superficial epidermal cells **(Supp Fig 14)**^31^. In the intestine, multipotent *Lgr5*+ stem cells in crypt base display elevated steady-state and nascent transcripts compared to the more terminally differentiated enterocyte and brush cell populations **(Supp Fig 15A-E)**. These stem cells, which rapidly replenish the colonic epithelium during homeostatic turnover, widely display the previously explored hallmarks of hypertranscription **(Supp Fig 15F-H)**^47^. Independent support for our findings is provided by Jao and Salic, who reported that intestinal crypts, where stem/progenitor cells reside, have high levels of incorporation of EU, a marker of nascent transcription^34^. Interestingly, we also observed a substantial enrichment of transcripts in goblet cells, which are required for continuous mucin production **(Supp Fig 15D)**. The presence of high transcript counts in this population adds further support to a novel role of hypertranscription in driving secretory cell output and is deserving of further investigation.

### Dynamic Modulation of Hypertranscription During Adult Cell Differentiation

We next looked to characterize dynamic changes in hypertranscription through the course of differentiation. We performed pseudotemporal ordering using Monocle on isolated hematopoietic cell trajectories representing the differentiation pathways of granulocytes, erythroblasts, and monocytes^48^. In granulocytes, we found that transcript abundance follows a gradual decrease throughout differentiation towards terminal identities **(Fig 4A-C)**. To further probe transcriptional dynamics during this differentiation path, we used k-means clustering to organize differentially expressed genes into 6 classes **(Fig 4D-I)**. Despite some overlap, these classes are enriched with unique biological processes, such as translational components in Class 1 and cell cycle genes in Class 5. Remarkably, we found that only Class 1 genes are downregulated substantially faster than the average total downregulation of transcriptional output over pseudotime, while all other classes track with the average or lag behind (colored lines in **Fig 4D-I)**. These results suggest that in the context of differentiation, the exit from hypertranscription may be primarily initiated via the downregulation of ribosome biogenesis and translational output. The preferential protein instability of euchromatin and transcriptional activators that we have previously reported in embryonic stem cells may similarly in adult cells make hypertranscription dependent on high translational capacity^10^.

**Figure 4:**
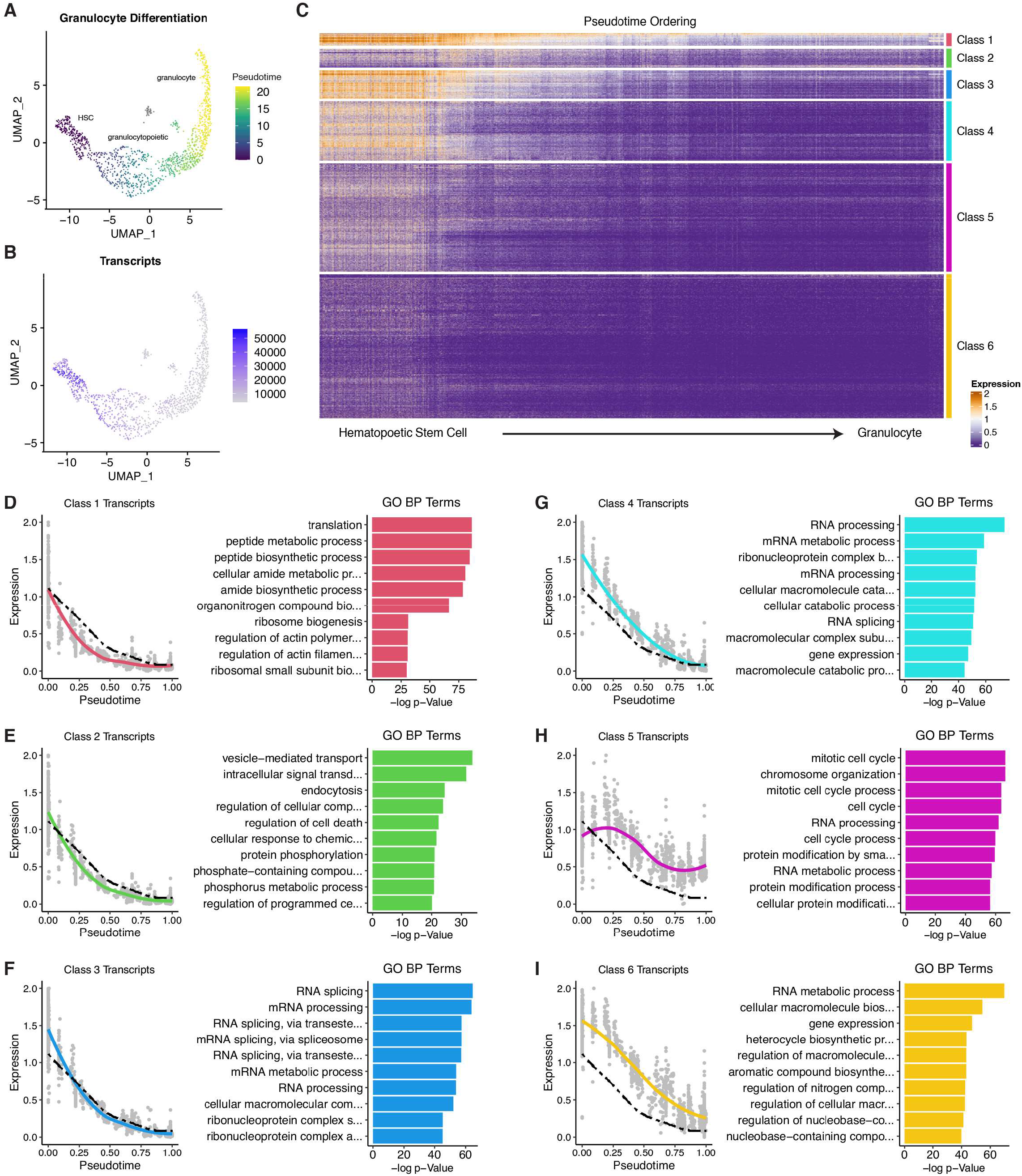
Hypertranscription Marks Developmental Progress in Hematopoiesis. **(A-B)** UMAP visualization of Monocle pseudotime scores and total transcript abundance under absolute scaling. Cd34+/cKit+ hematopoietic stem cells were defined as root cells for trajectory inference analysis. **(C)** Heatmap depicting transcriptomic expression changes across differentiation pseudotime from hematopoietic stem cell to granulocyte. Rows depict individual genes and columns depict single-cell transcriptomes ordered by pseudotime values. Class annotations represent k-means clusters with expression presented as z-scores. **(D-I)** Transcriptome curves representing expression of clustered genes across progression of pseudotime. Dotted line represents average expression of all genes (as depicted in **(C)**). Right panels of each column show enriched GO terms derived from genes within each class. Expression in **(C-I)** represents ranged log2 expression of ERCC-normalized reads.

In the erythroblast and monocyte trajectories, we observed similar results with progressive losses of total transcript content with pseudotime **(Supp Fig 16)**. In addition to this overall decline, we detected small populations of intermediate cells with transient increases in transcript content, reflected most strikingly in the levels of ribosomal and housekeeping genes **(Supp Fig 16J, K)**. Interestingly, these brief bursts of increased transcription coincide with the onset of key differentiation effector genes that rapidly rise to very high levels, such as the induction of beta-2 hemoglobin in developing erythroblasts **(Supp Fig 16N)**. It is possible that these transient rises in transcriptional output support high levels of synthesis needed for the rapid production of differentiation effector proteins. Importantly, while global scaling would correctly identify the onset of these differentiation markers, it would entirely miss the remarkable dynamics of global transcription during lineage commitment, including large scale shifts in the levels of the transcription and translation machineries **(Supp Fig 16O)**. Taken together, our analyses uncover a remarkably rich level of dynamics in global transcription at the single cell level and point to a pervasive redeployment of hypertranscription beyond embryogenesis in adult stem/progenitor cells.

### Rapid Induction of Hypertranscription During Adult Organ Regeneration

The observation that hypertranscription is dynamically regulated during physiological renewal of adult organs led us to explore its status during regeneration. We first focused on the context of skeletal muscle repair, which is mediated by the rapid proliferation and differentiation of resident muscle satellite stem cells^49^. This context was of particular interest as steady-state satellite stem cells display relatively low transcript abundance in the Tabula Muris (see **Fig 2, Table S1**). Remarkably, we found that the total transcript content of satellite stem cells ranks highest compared to other cell types across a dataset of cardiotoxin-induced tibialis muscle injury^50^ **(Fig 5A-C)**. When stratified into specific days post-injury (DPI), we found that non-injured and 21 DPI satellite stem cells display comparably low transcript content **(Fig 5D)**. In contrast, 0.5 DPI satellite stem cells display a notable 3.5-fold upregulation of transcriptional output, which peaks at 2 DPI and slowly returns to baseline over the next 19 days. This rapid elevation of transcription upon injury precedes a large increase in the numbers of satellite stem cells, which rise by nearly two orders of magnitude across a 3-day time period **(Fig 5D)**. In agreement with our data on hypertranscribing hematopoietic and epithelial stem/progenitor cells **(Figs 2-4, Supp Fig 14-15)**, we found that the genes most differentially expressed in 0.5 DPI vs non-injured satellite stem cells are enriched for functions in ribosome biogenesis, translation regulation and RNA processing **(Fig 5E)**. Together, these data indicate that muscle satellite stem cells enter a state of injury-induced hypertranscription that precedes rapid expansion and subsequent regeneration.

**Figure 5:**
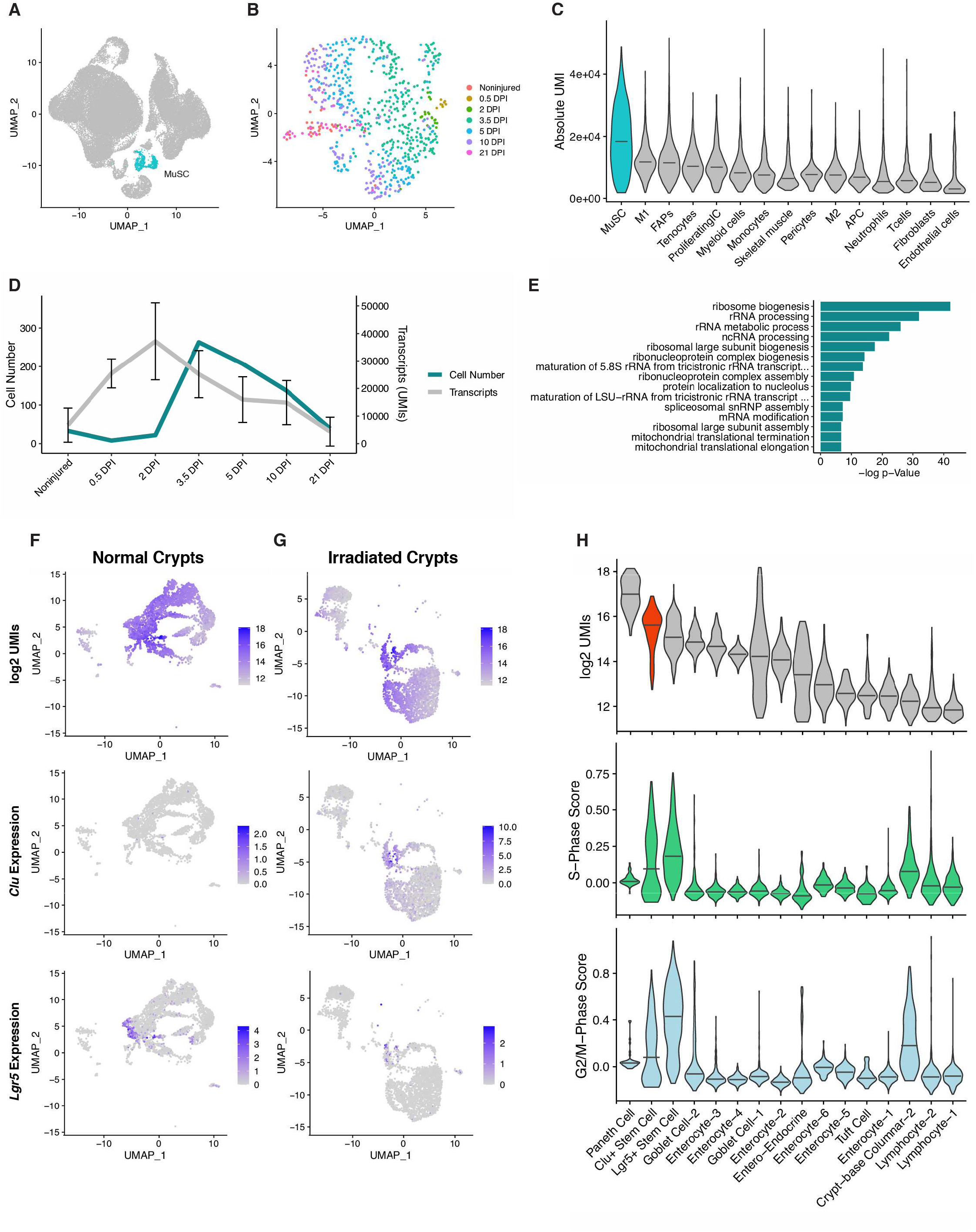
Hypertranscription is Deployed During Regenerative Processes. **(A)** UMAP visualization of all cells in the muscle repair atlas. Highlighted cells represent annotated satellite cells across all time points. Plot depicts dimensionality reduction performed under absolute scaling. **(B)** UMAP visualization of muscle satellite cells. Plot depicts dimensionality reduction performed under absolute scaling. **(C)** Distribution of transcript content between cell types across all time points. Lines correspond to median values. **(D)** Progression of satellite cell transcript content and cell number through injury and regeneration. Transcript counts represent raw UMIs, error bars represent standard deviation. **(E)** GO terms enriched in >1.5-fold differentially upregulated genes in 0.5 DPI vs non-injured satellite cells. **(F-G)** UMAP visualization of transcript abundance, stem cell marker *Lgr5*, and revival stem cell marker *Clu* across all cells in normal/irradiated crypts. Plots depict dimensionality reduction performed under absolute scaling. **(H)** Distribution of transcript abundance and cell-phase scores between cell types in normal and irradiated crypts. Cell types are ranked by median log2 UMIs with lines corresponding to median values.

In a separate model of organ damage, we explored the recovery of the intestinal epithelium to irradiation. Canonical intestinal stem cells, marked by Lgr5 expression, reside at the base of intestinal crypts and renew the epithelium during normal homeostasis^51^. However, recent work has demonstrated that a rare, separate type of cell, named revival stem cells and marked by Clu expression, are responsible for regeneration in response to irradiation^52^. Consistent with our analysis in the Tabula Muris (see **Fig 2**), Lgr5+ intestinal stem cells display the highest transcript content amongst all cell types in the crypt compartment **(Fig 5F-H)**. Interestingly, upon irradiation-induced loss of Lgr5+ intestinal stem cells, Clu+ revival stem cells assume comparably high levels of transcript content and cell cycling **(Fig 5H)**. These findings suggest that, as Clu+ revival stem cells exit quiescence to replenish the crypt niche, they enter hypertranscription to support the biosynthetic requirements of regenerative proliferation and differentiation. Thus, our results indicate that hypertranscription is redeployed in adult stem/progenitor cells both during physiological organ renewal as well as in different contexts of regeneration upon injury.

## DISCUSSION

We report here the development and validation of absolute-scaling approaches to estimate transcript abundance in scRNA-seq data. When applied to a variety of scRNA-seq datasets, absolute scaling accurately captures known cases of hypertranscription in embryonic cells and uncovers a previously unappreciated rich level of global transcriptional dynamics in adult cells. Our analyses indicate that hypertranscription is pervasively redeployed by stem/progenitor cells across adult organ systems under both homeostatic and regenerative conditions. Thus, rather than being largely ignored as it is by standard scRNA-seq analyses methods, we propose that hypertranscription is central to the biology of adult organs and needs to be considered in order to understand the mechanisms that underlie their maintenance and regeneration.

Our results document that absolute scaling is capable of capturing large-scale changes in transcriptional output in scRNA-seq data, as long as ERCC exogenous spike-ins or UMI barcodes are used. While both methods remain useful, our data slightly favor the use of ERCCs over UMIs in this context, likely because they are introduced further upstream in sample processing. Nevertheless, several potential sources of technical variation remain. ERCC-based analysis is dependant on proper experimental aliquoting, and whether various technical biases equally affect endogenous and extrinsic sequences remains debated^14^. The accuracy of UMIs, in addition to aforementioned issues, can also be affected by fluctuations in sequencing depth^14^. As with all scRNA-seq studies, the use of higher cell numbers and greater read depths per cell attenuate sources of potential artifactual noise. Nonetheless, our ground truth tests and reproduction of embryonic hypertranscription results strongly validate the ability of absolute scaling to detect hypertranscription and its associated features in scRNA-seq data. Our results underscore the notion that global variations in total cellular mRNA content represent an important but critically underappreciated dimension in many biological contexts. The capacity to explore hypertranscription in single-cell data will enrich our understanding of many biological questions, notably in adult organ maintenance, regeneration, pathology and malignancy. We anticipate that this will also be aided by the development of improved protocols that more quantitatively capture the composition of total cellular RNA, including ribosomal and non-coding RNA species. Importantly, global scaling and absolute scaling are not mutually exclusive and can be applied to the same dataset, maximizing the information that is extracted from each experiment. Moreover, current methods for inference of cell trajectories in scRNA-seq data like pseudotime align the cells by transcriptome similarity but do not systematically annotate the start/end points of the trajectories, instead usually relying on manual annotation^53^. We suggest that hypertranscription may be used to anchor the start of such trajectories, particularly in contexts of cell differentiation. In addition, hypertranscription provides a new dimension with which to refine heterogeneous cell populations that may express similar markers. This may be particularly useful to distinguish quiescent vs activated stem cells, as we observed for satellite cells in the muscle regeneration paradigm above **(Fig. 5A-E)**.

The existence of high transcript-content cells occupying adult stem cell and progenitor niches expands the role of hypertranscription beyond development and tumorigenesis. Our findings are supported by early studies dating back to the 1930’s, reporting evidence of elevated total RNA content within progenitor cells of the hematopoietic and intestinal systems^54–56^. In parallel, our findings of hypertranscription in the context of regeneration are closely aligned with classical studies showing increased RNA content in renewing cells of planarians and salamanders^57,58^. Our work builds upon these studies by comprehensively exploring transcriptional heterogeneity and dynamics at the single-cell and transcriptome-wide levels. In the process, our analyses uncover putative molecular drivers of hypertranscription in diverse contexts of adult organ homeostasis that warrant further studies.

We identify hypertranscription as a feature of active stem/progenitor cells that decays along lineage commitment. We note that the transcriptome is not linearly amplified or repressed, but follows patterns such as the primacy of modulation of regulators of ribogenesis and translation that are not fully understood and deserve further investigation^9,10^. Moreover, transcript content does not decline strictly linearly down differentiation pathways but rather includes transient fluctuations, possibly due to high synthesis of differentiation effector proteins or expansion of transit amplifying-like cells (see **Supp Fig 16**). Further dissection of these transcriptional dynamics is likely to reveal new aspects of the biology of stem cell compartments. The existence of high transcript-content cells occupying adult stem cell and progenitor niches greatly expands the prevalence of hypertranscription beyond development and tumorigenesis^2,13^. Our initial analyses reveal many similarities between adult hypertranscription and its embryonic counterpart, but there are likely to be adult-specific and/or organ-specific aspects of the regulation of hypertranscription. It will be of interest to identify the primary drivers of the onset of and exit from hypertranscription in different organs, decipher the chromatin dynamics that underlie these changes, and dissect how they integrate with signaling from the stem/progenitor cell niche, both during physiological organ homeostasis and regeneration after injury. Our work lays a foundation for the investigation of hypertranscription in adult organs at the single cell level.

Early histochemical studies dating back to the 1950’s reported evidence of elevated total RNA and protein content within renewing cells of planarians and salamanders, but these findings have essentially been forgotten in the genomics era^57,58^. Our results put a new focus on hypertranscription and its molecular hallmarks that can now be applied to a plethora of regeneration paradigms being explored with scRNA-seq approaches, such as axolotl limb or mouse digit tip regeneration^59–62^. Of note, it has recently been shown that during generation of iPS cells a combination of high proliferative capacity with hypertranscription identifies cells with the greatest probability to faithfully reprogram to pluripotency^63^. We anticipate that further exploration of hypertranscription in the context of regeneration and cellular reprogramming paradigms will provide fundamental new molecular insights that may have broad impact in regenerative medicine.

## METHODS

### Data Sources

Datasets and software versions used for this study are outlined in **Supp Fig 1C**. Raw read data from published studies were downloaded from either ENA or SRA. These include ArrayExpress accession code E-MTAB-2600^25^ and Gene Expression Omnibus accession codes GSE118767^23^, GSE96583^21^, GSE108313^20^, GSE138826^50^, GSE136441^26^, GSE109774^31^, and GSE123516^52^. The 12K 1:1 HEK293T-NIH3T3 mixture dataset was obtained from 10X Genomics under the Chromium v2 Chemistry Demonstration.

### Pre-Processing of scRNA-seq Data

Detailed parameters for analysis and figure scripts are provided at https://github.com/yunkyokim/AbsoluteScaling. Published count matrices with collapsed UMIs or mapped ERCC sequences were used where available. For CEL/Sort-seq data from Tian et al 2019, new count matrices were generated using the scPipe pipeline using genomic references containing ERCC sequence data^64^. Spliced/unspliced count matrices for the bone marrow, intestine, and skin FACS datasets were generated using the scVelo pipeline for analysis of nascent reads^65^.

Seurat objects were used for processing of all scRNA-seq data. Low-quality cells in individual organ datasets were filtered by mitochondrial gene expression, ERCC fraction, and gene/transcript counts. To address the possibility that high-content cells were the result of libraries generated from doublets, we performed heterotypic doublet removal on Smartseq2 and 10X Genomics datasets using DoubletFinder^32^. The total doublet rate was estimated at 4%, and the homotypic doublet rate was derived from Tabula Muris cell-type annotations. All doublet removal was performed on globally-scaled data with standard processing.

### Scaling and Dimensional Reduction

Absolute scaling analysis of ERCC datasets was performed using the scran package^66^. To ensure that transcript counts between organs were comparable, individual organ count matrices were merged and per-cell size factors were generated from spike-in data using the computeSpikeFactors function. This calculation is derived from the sum of all spike-in reads and generates a set of size factors with a mean of unity. Single-cell libraries were then normalized to size factors and log2 scaled. For absolute scaling analysis of UMI datasets, raw UMI counts were used following log2 scaling. Global scaling was performed on ERCC and UMI datasets by first removing spike-in data (if present) and normalizing for library size using the relative counts methods in Seurat’s NormalizeData function. Results were subsequently log2 scaled and used for analysis.

Standard dimensional reduction was performed using Seurat functions with detailed parameters available in code. Briefly, highly variable features were scaled and used for principal component analysis, with the number of components determined using resampling tests^67^. Data was subsequently visualized using the Uniform Approximation and Projection Method (UMAP).

### Cell Cycle Scoring and Signature Analysis

To determine cell phase, we used the CellCycleScoring method from Seurat with markers as previously published^35^. Briefly, individual cells were scored based on the expression of either G2/M or S phase cell markers. Cells lacking expression of either marker set were identified to be in a non-cycling G1 phase. For signature analysis, we used the VISION tool to calculate signature scores using gene sets derived from the Molecular Signatures Database^68,69^. Signatures were calculated using absolute-scaled data without internal transformation within VISION. For the “Serum Hypertranscription” signature, >2.5-fold differentially upregulated genes in serum mESCs (Kolodziejczyk et al 2015) were used to generate the gene set.

### Quantile Polarization Scores and Pseudotemporal Ordering

Quantile polarization scores were calculated with VECTOR using the vector.getValue function. As previously described, we determined quantile polarization values on absolute-scaled data using the top 150 principal components^44^. For pseudotemporal analysis with Monocle 3, we converted pre-processed and absolute-scaled Seurat objects into Monocle3 cds objects. Cells were clustered on previously generated UMAP reductions using the Leiden algorithm. To construct single-cell trajectories, hematopoietic stem cells expressing Cd34 and Kit were used as root cells. Principal graphs were constructed and used for pseudotemporal ordering.

### Quantification and Statistical Analysis

All statistical analysis was performed using functions implemented in R. Spearman correlation was used to identify signatures related to cellular transcript abundance and was calculated using cor. Transcriptome curves were drawn using geom smooth in ggplot2 using a “loess” model. Cumulative distributions plots for mESC transcriptomes were generated using ecdf. For differential gene expression in scRNA-seq data, we used the FindMarkers functions with Seurat and performed gene ontology and ChEA enrichment analysis in Enrichr^70^.

## Supporting information

Table S1 - Tabula Muris Absolute Scaling Stats

Table S2 - Gene-Transcript Content Correlations

Table S3 - GO BP and ChEA Terms Enriched in Highly Correlated Genes

## ACKNOWLEDGMENTS

We thank Gary Bader, Aydan Bulut-Karslioglu, Kieran Campbell, Joshua Currie, Michelle Percharde, and members of the Santos Lab for input and critical reading of the manuscript. This work is supported by a Whiteside Scholarship (University of Toronto) and a McLaughlin Award (University of Toronto) to Y.K., CIHR Project Grant 420231, and a Canada 150 Research Chair in Developmental Epigenetics to M.R.-S.

## AUTHOR CONTRIBUTIONS

Y.K. and M.R-S. conceived of the project. Y.K. performed all analyses. M.R-S. supervised the project. Y.K. and M.R.-S. wrote the manuscript.

## SUPPLEMENTARY FIGURE LEGENDS

**Supplementary Figure 1:**
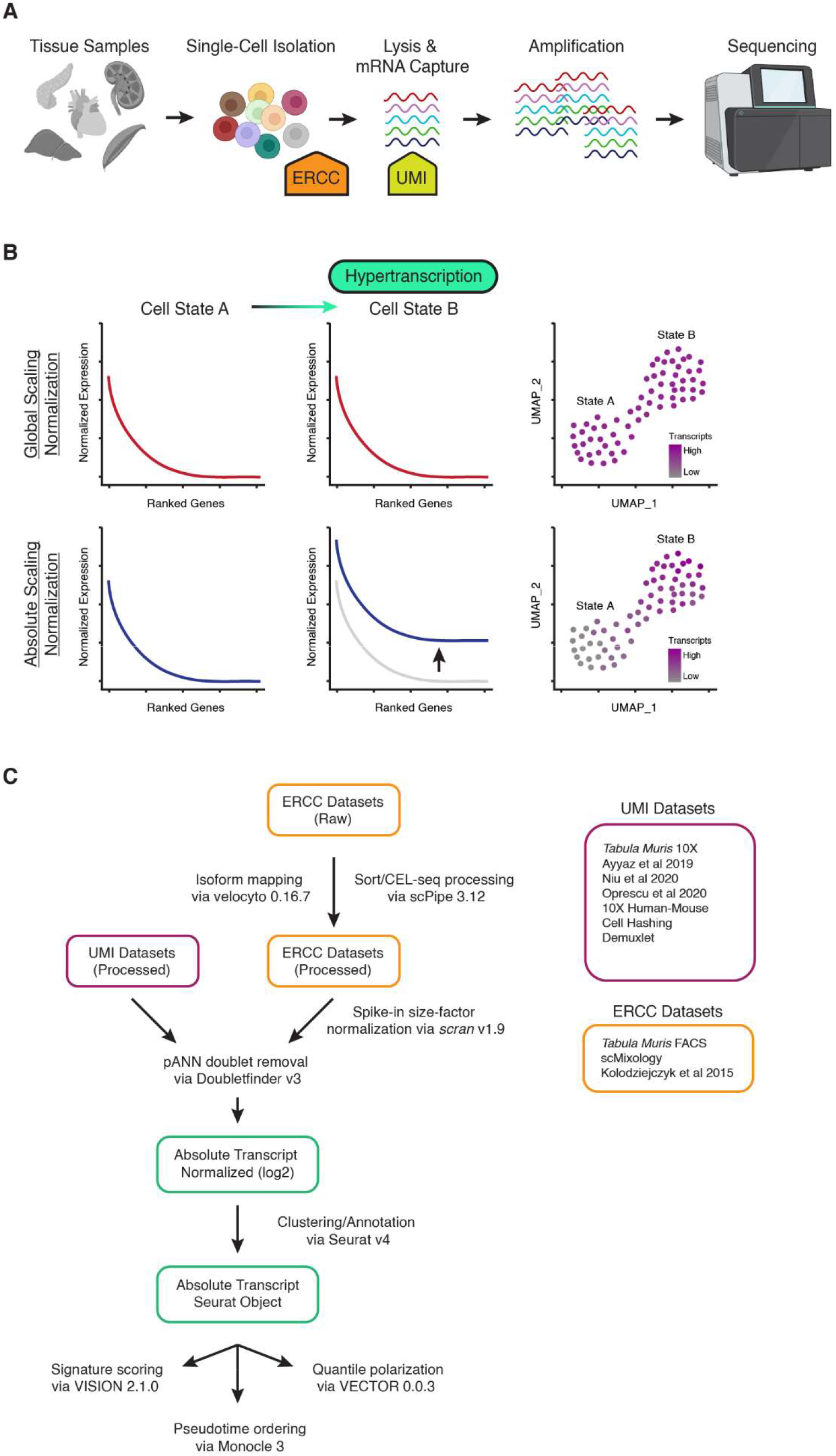
Absolute Scaling of Single-Cell Datasets. **(A)** Schematic of ERCC and UMI tools within scRNA-seq protocols. Tissue samples are disassociated into single cells and lysed to capture mRNA transcripts. ERCC spike-in sequences are added to lysed contents and captured alongside endogenous mRNA species. UMIs are added during the capture and RT steps to uniquely label transcripts. **(B)** Model of hypertranscription in scRNA-seq datasets. Absolute scaling preserves large-scale transcriptomic shifts that are typically masked in global scaling approaches due to assumptions of mRNA content parity between cells of a dataset. **(C)** Summary of bioinformatic tools and datasets used for absolute scaling in this study.

**Supplementary Figure 2:**
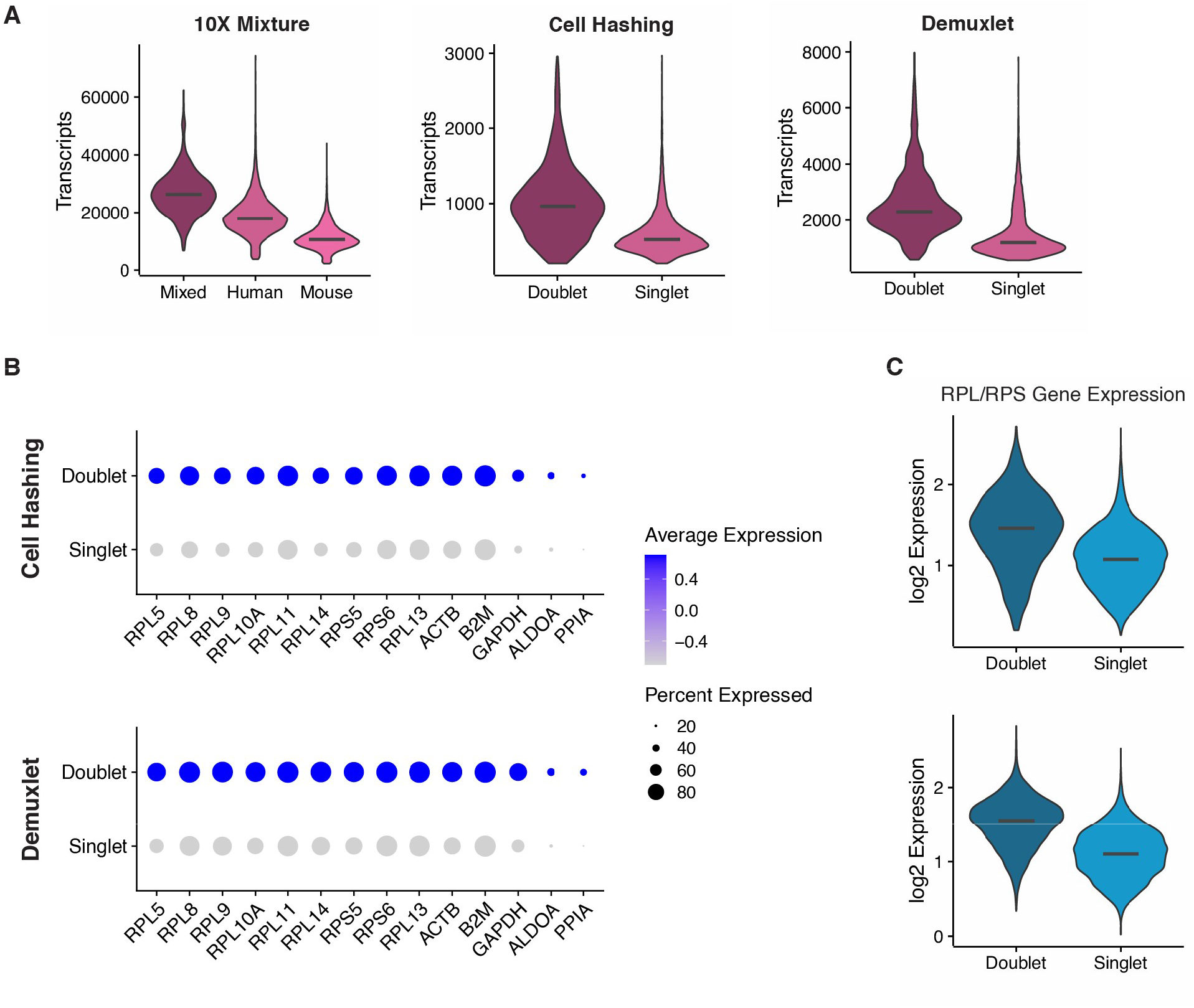
Absolute Scaling Reproduces Transcript Content Differences Between Singlets and Doublets. **(A)** Transcript abundances in singlets and doublets within ground truth datasets. Transcripts represent raw UMIs, lines correspond to median values. **(B)** Dot plots depicting expression of ribosomal/housekeeping genes under absolute scaling in PBMC ground truth datasets. Expression represents z-scores of log2-scaled raw UMIs. **(C)** Average expression of RPL and RPS gene expression in ground truth PBMC datasets. Lines correspond to median values.

**Supplementary Figure 3:**
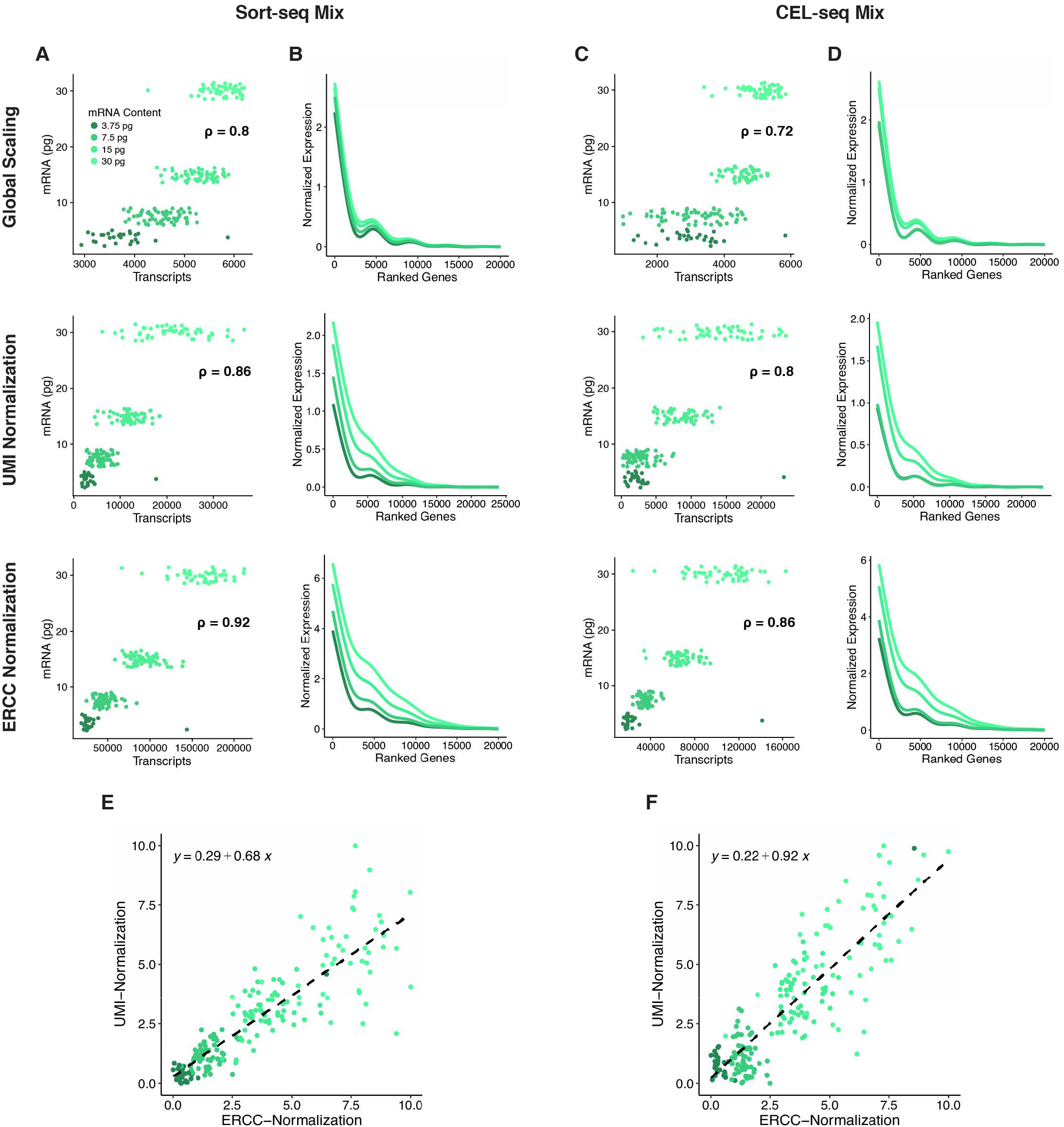
Absolute Scaling Reproduces Pseudocell Transcript Content in Benchmarking Mixtures. **(A, C)** Transcript abundances per pseudocell vs ground truth mRNA content under different normalization methods. Each point represents a single artificially-generated pseudo-cell transcriptome using defined amounts of cell-line mRNA. **(B, D)** Transcriptome curves depicting gene expression across top 25,000 genes in pseudocells. Individual genes are ranked using combined log2 expression between all pseudocell ground truth conditions. **(E, F)** Comparison of UMI vs ERCC absolute scaling. Ranged expression represents log2 raw UMIs or ERCC-normalized reads.

**Supplementary Figure 4:**
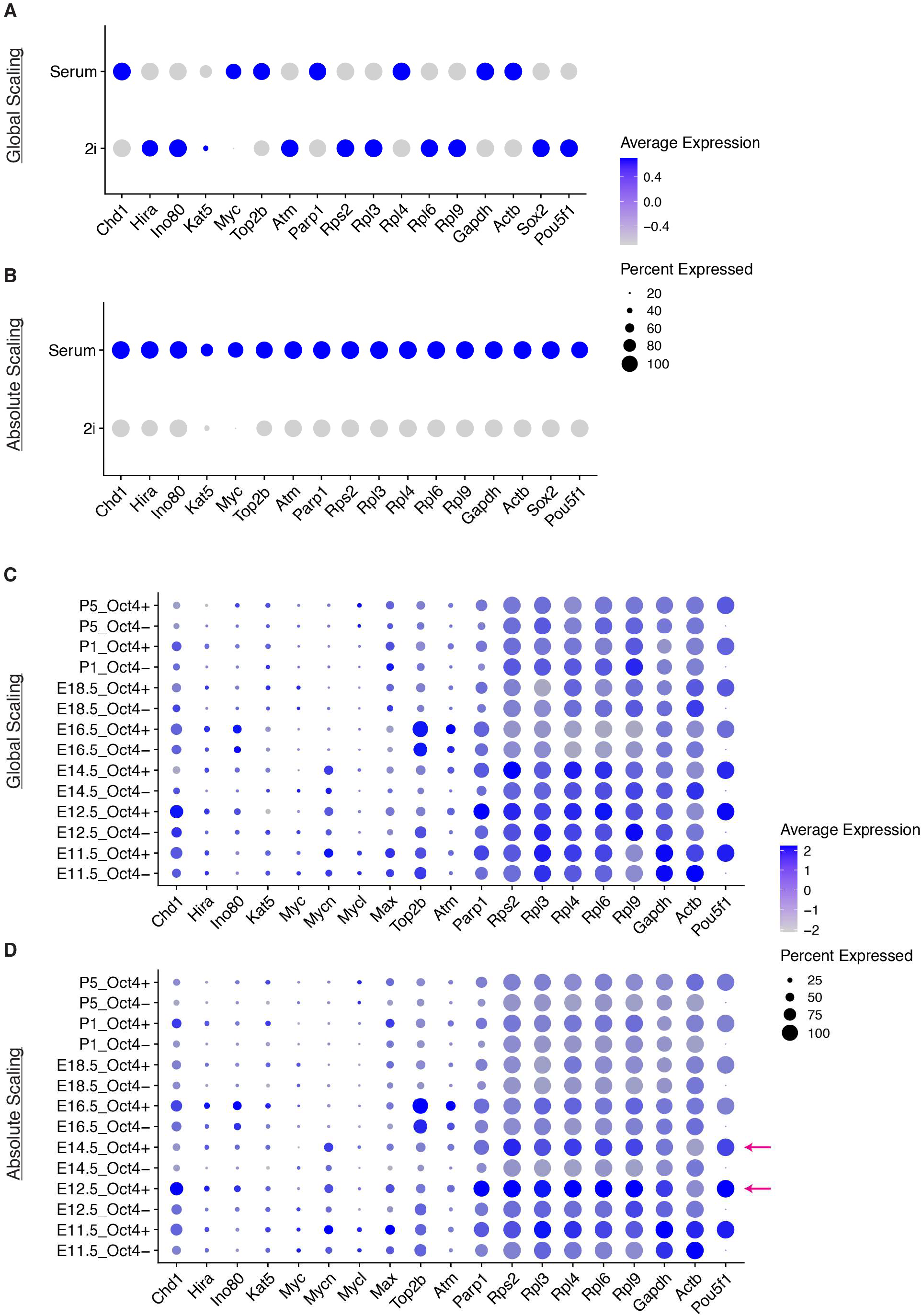
Absolute Scaling Captures Hypertranscription Hallmarks in Embryonic Datasets. **(A-B)** Expression of genes relevant to hypertranscription hallmarks in serum/2i mESCs under global vs absolute scaling. **(C-D)** Expression of genes relevant to hypertranscription hallmarks in Oct4+ PGCs/Oct4- soma under global vs absolute scaling. Expression represents z-scores of log2-scaled transcripts.

**Supplementary Figure 5:**
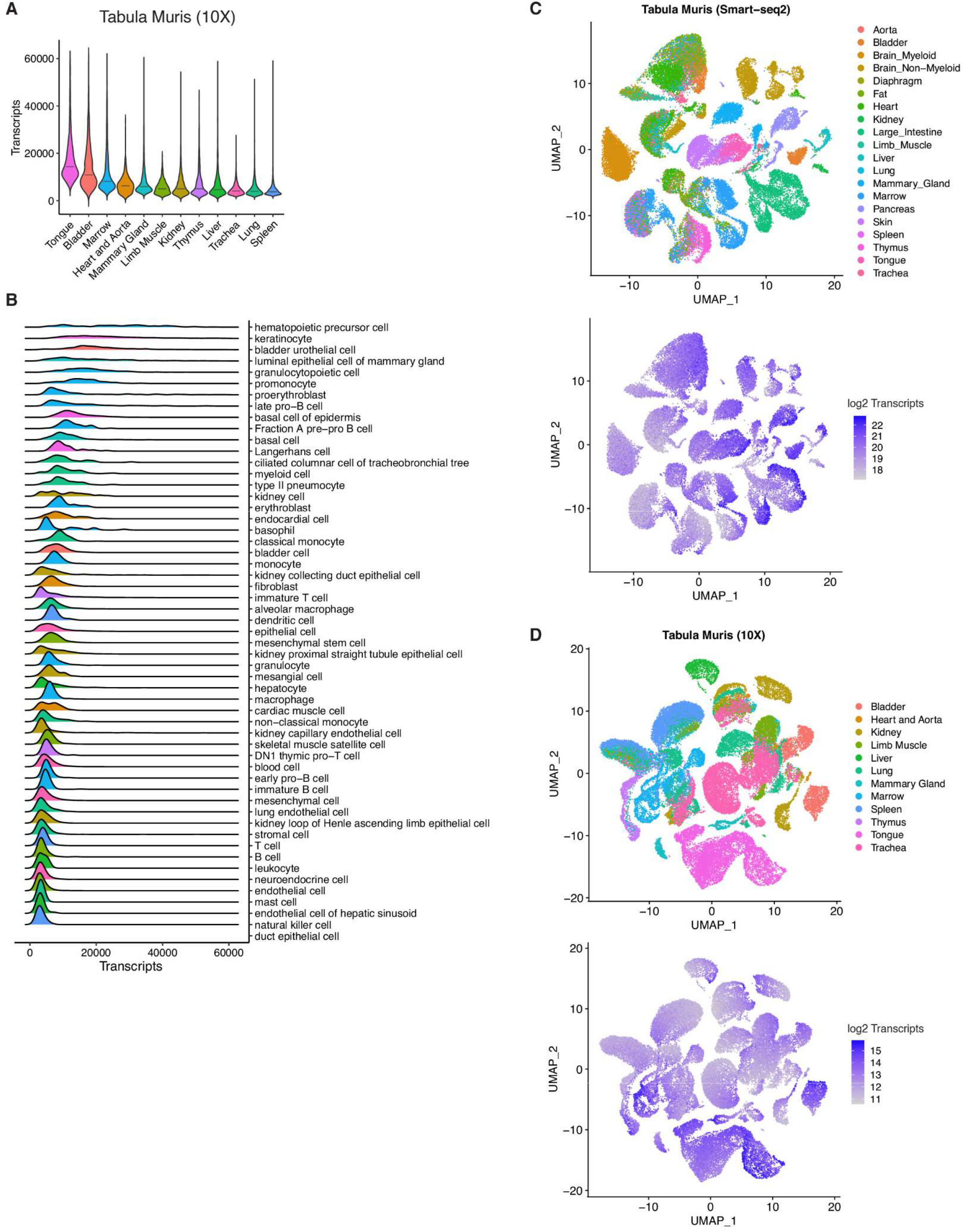
Transcript Content Heterogeneity in Tabula Muris 10X Datasets. **(A-B)** Distribution of cellular transcript content between Droplet organ datasets under absolute scaling, ranked by median total transcripts. Transcripts represent raw UMIs, lines correspond to median values. **(C-D)** UMAP representations of FACS/Droplet Tabula Muris Atlas generated under globally-scaled gene expression. Transcript abundance represents log2 scaled ERCC-normalized reads **(C)** or log2 absolute UMIs **(D)**. Plots depict dimensionality reduction performed under absolute scaling.

**Supplementary Figure 6:**
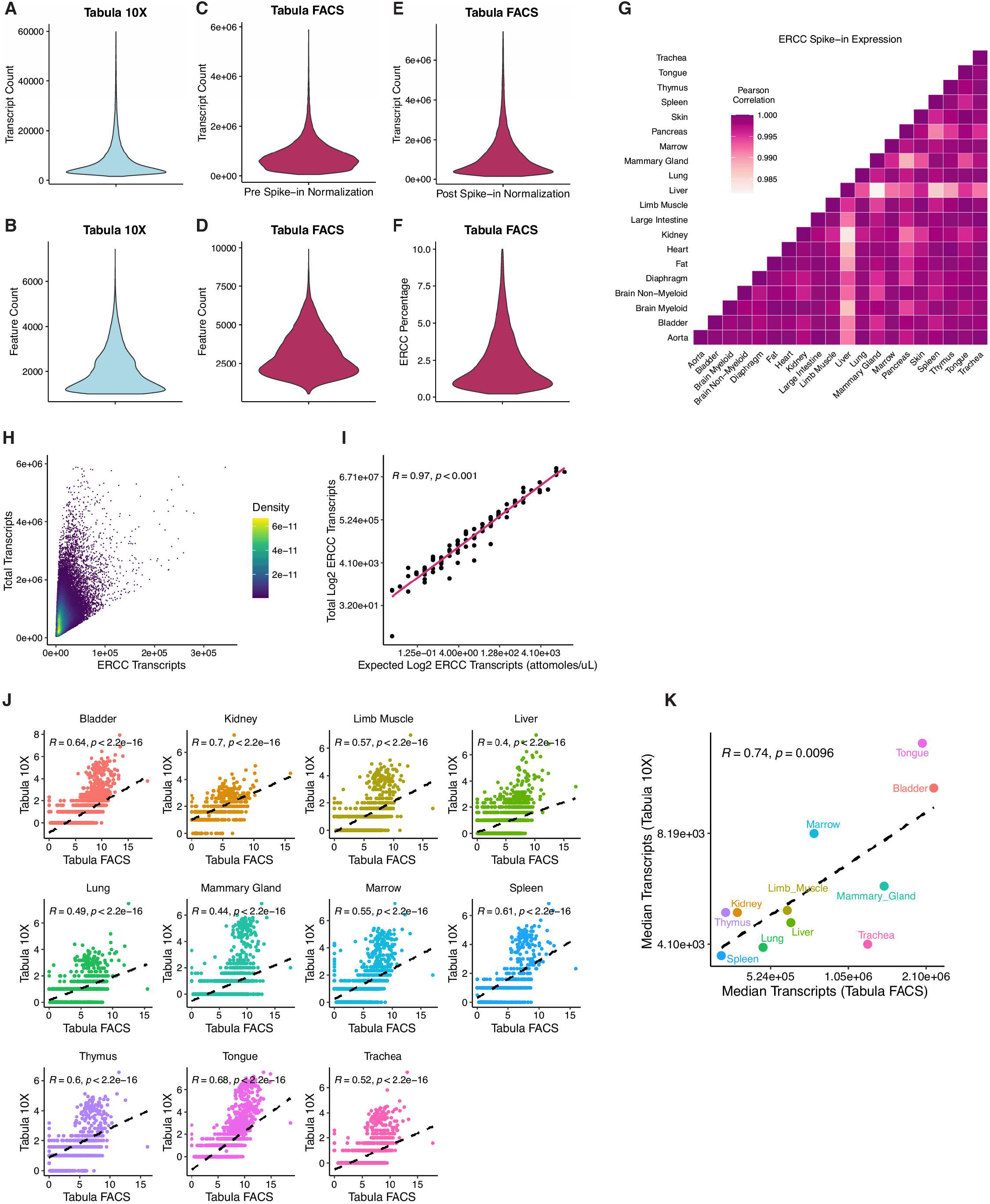
Quality Control of Tabula Muris. **(A-F)** Transcript and feature counts of Tabula Muris FACS or Droplet atlases. **(G)** Linear correlations of ERCC spike-in expression between FACS organ datasets. Values represent Pearson coefficients. **(H)** Representation of filtering in FACS datasets for transcript counts and ERCC percentage. **(I)** Comparison of actual and expected ERCC species expression based on spike-in mix composition. **(J)** Comparison of individual gene expression between FACS and Droplet datasets. Each point represents a single shared gene between FACS and Droplet dataset. **(K)** Comparison of average transcript abundance in organ datasets between FACS and Droplet atlases. For **(J-K)**, transcripts in FACS represents log2 ERCC-normalized expression, while values in Droplet represent log2 absolute UMIs. Correlation values represent Pearson coefficients.

**Supplementary Figure 7:**
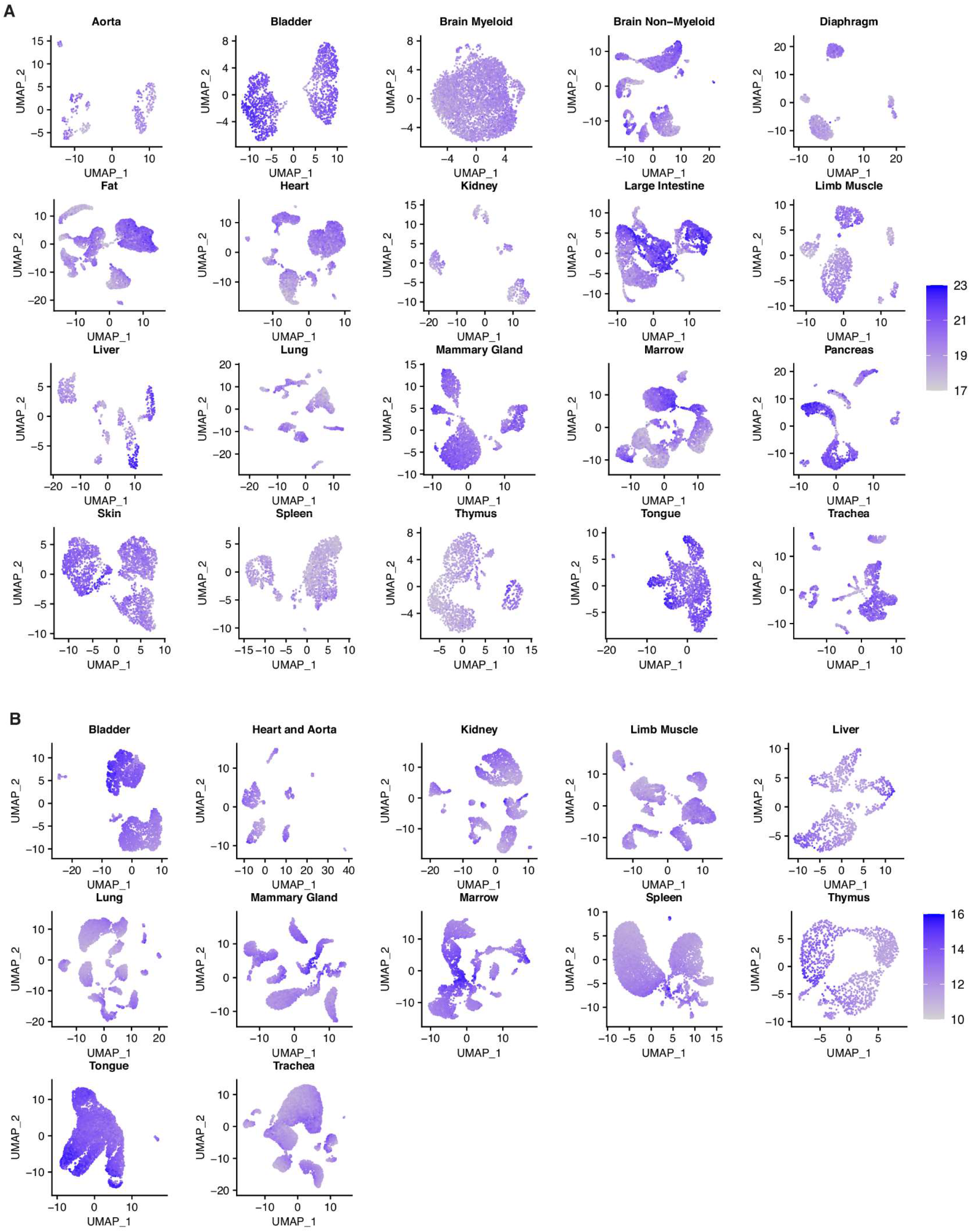
Absolute Scaling Reveals Clustering of High-Transcript Content Cells. **(A-B)** UMAP representations of FACS/Droplet organ datasets using dimensionality reduction under globally-scaled gene expression. Transcript abundance represents log2 scaled ERCC-normalized reads **(A)** or log2 absolute UMIs **(B)**.

**Supplementary Figure 8:**
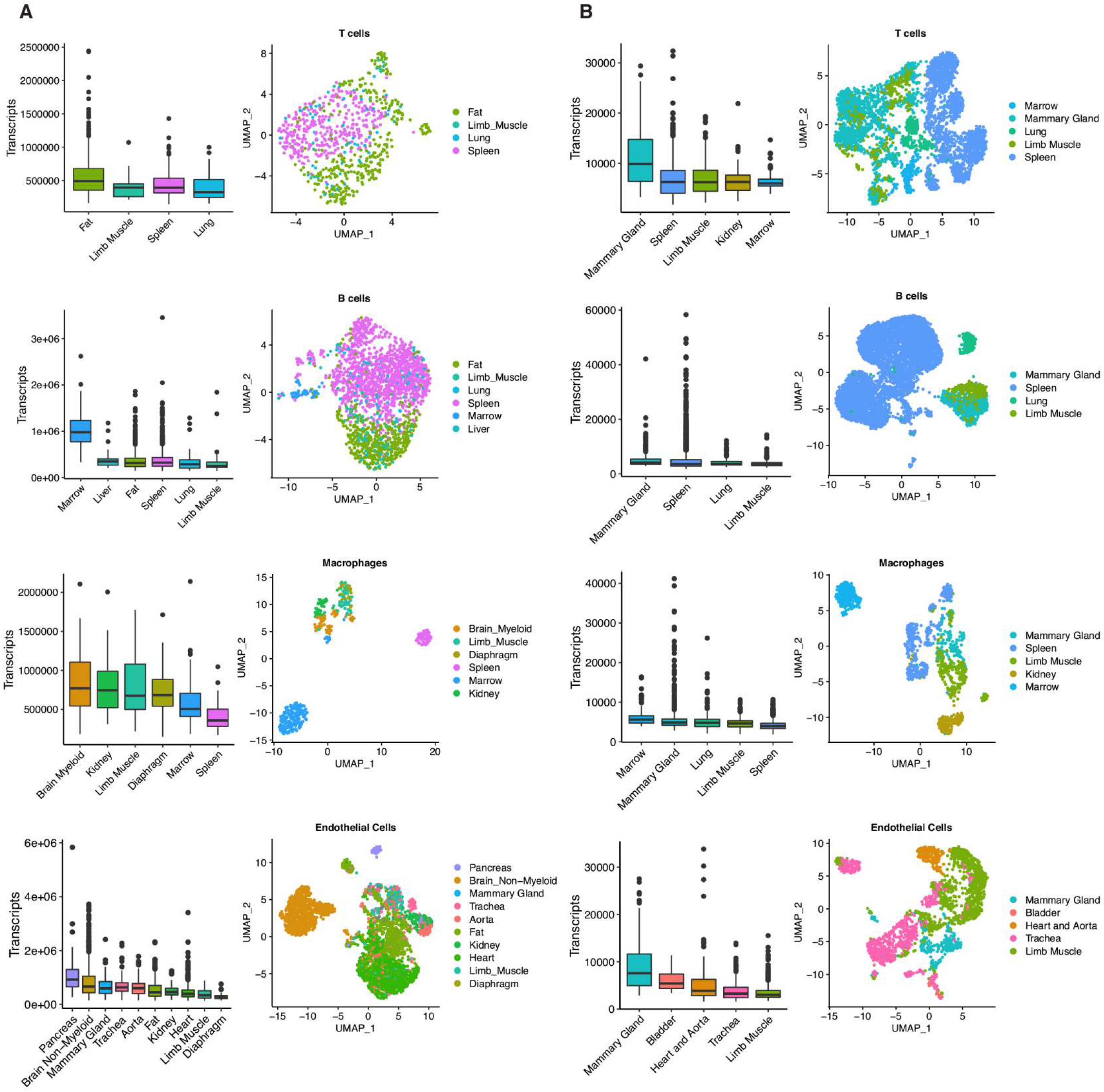
Shared Cell Types Between Organ Datasets Display Similar Transcript Abundances. **(A-B)** Boxplots of transcript content and UMAP visualizations of shared cell types in FACS **(A)** or Droplet **(B)** datasets. Cell types ranked by median transcript content. Values in **(A)** represent log2 scaled ERCC-normalized reads, while values in **(B)** represent log2 absolute UMIs.

**Supplementary Figure 9:**
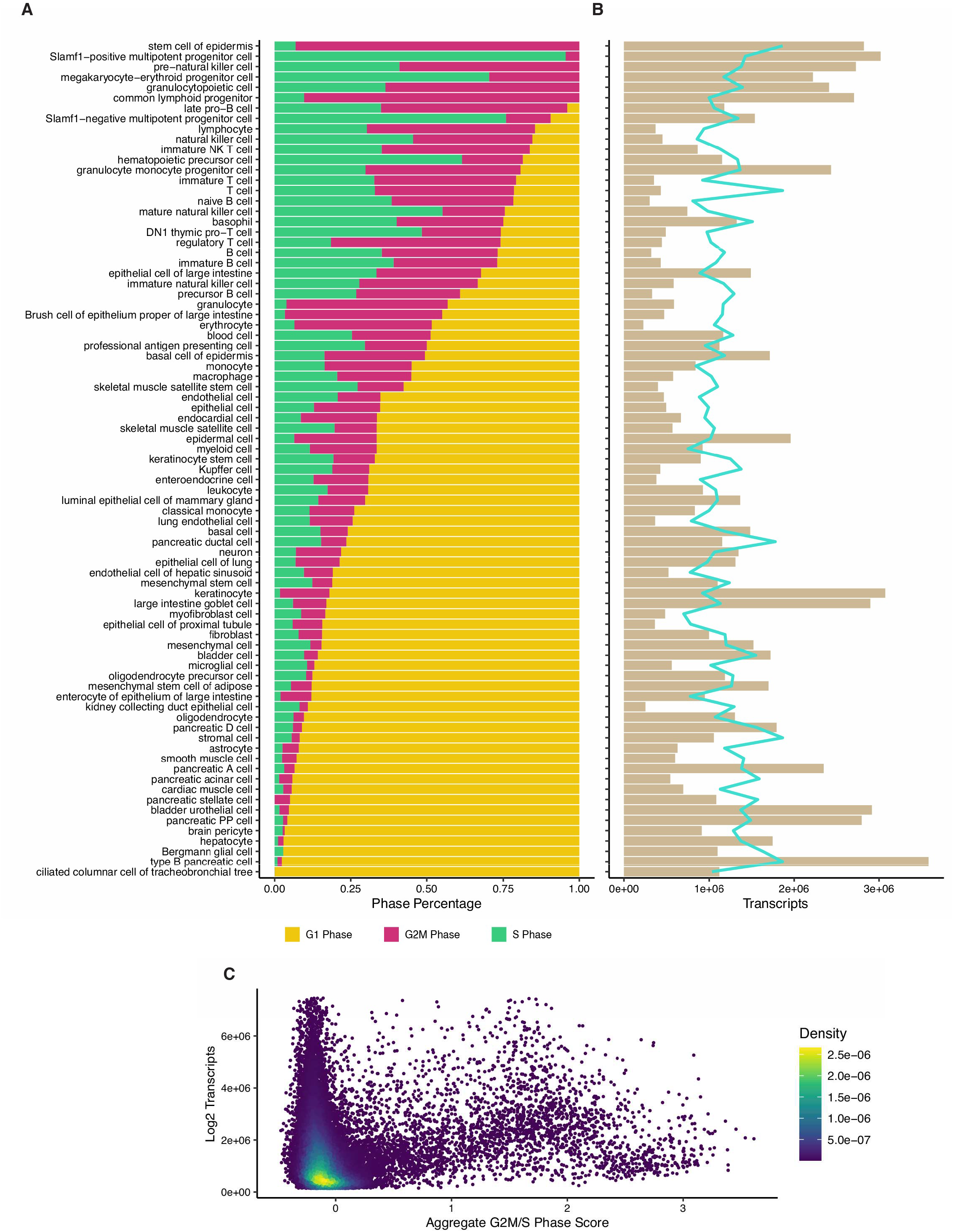
Cell Cycle Phase Distribution in Across FACS Atlas. **(A)** Cell cycle phase proportions across all cell types, sorted by percentage of G2/M and S cells. Phase scores were calculated using globally-scaled expression data. **(B)** Transcript abundance across all cell types, sorted by percentage of G2M and S cells. Transcripts represent log2 scaled ERCC-normalized reads. Line represents moving average across 7 cell types. **(C)** Comparison between aggregate G2M/S score and transcript abundance across all cells.

**Supplementary Figure 10:**
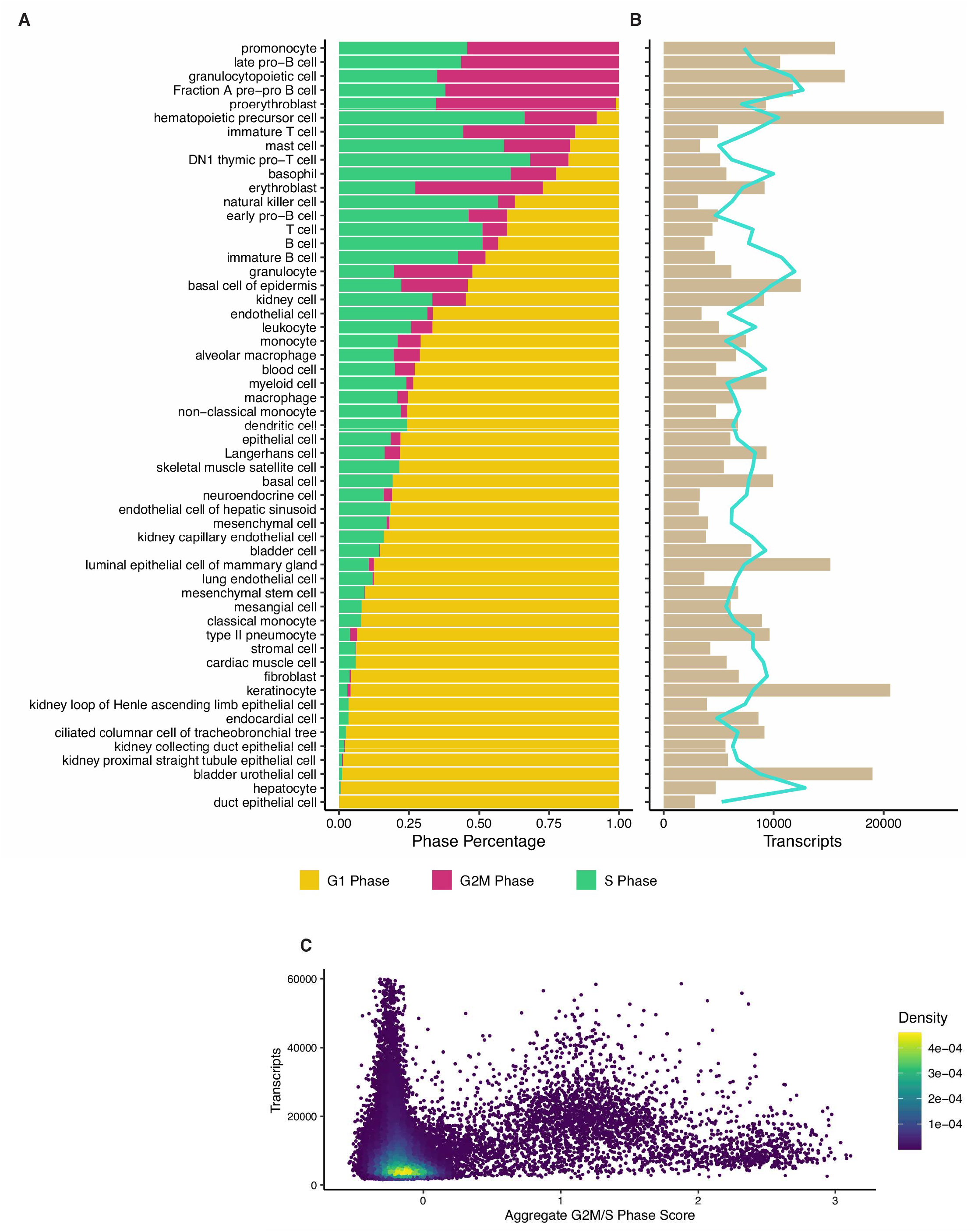
Cell Cycle Phase Distribution in Across Droplet Atlas. **(A)** Cell cycle phase proportions across all cell types, sorted by percentage of G2/M and S cells. Phase scores were calculated using globally-scaled expression data. **(B)** Transcript abundance across all cell types, sorted by percentage of G2M and S cells. Transcripts represent log2 absolute UMIs. Line represents moving average across 7 cell types. **(C)** Comparison between aggregate G2M/S score and transcript abundance across all cells.

**Supplementary Figure 11:**
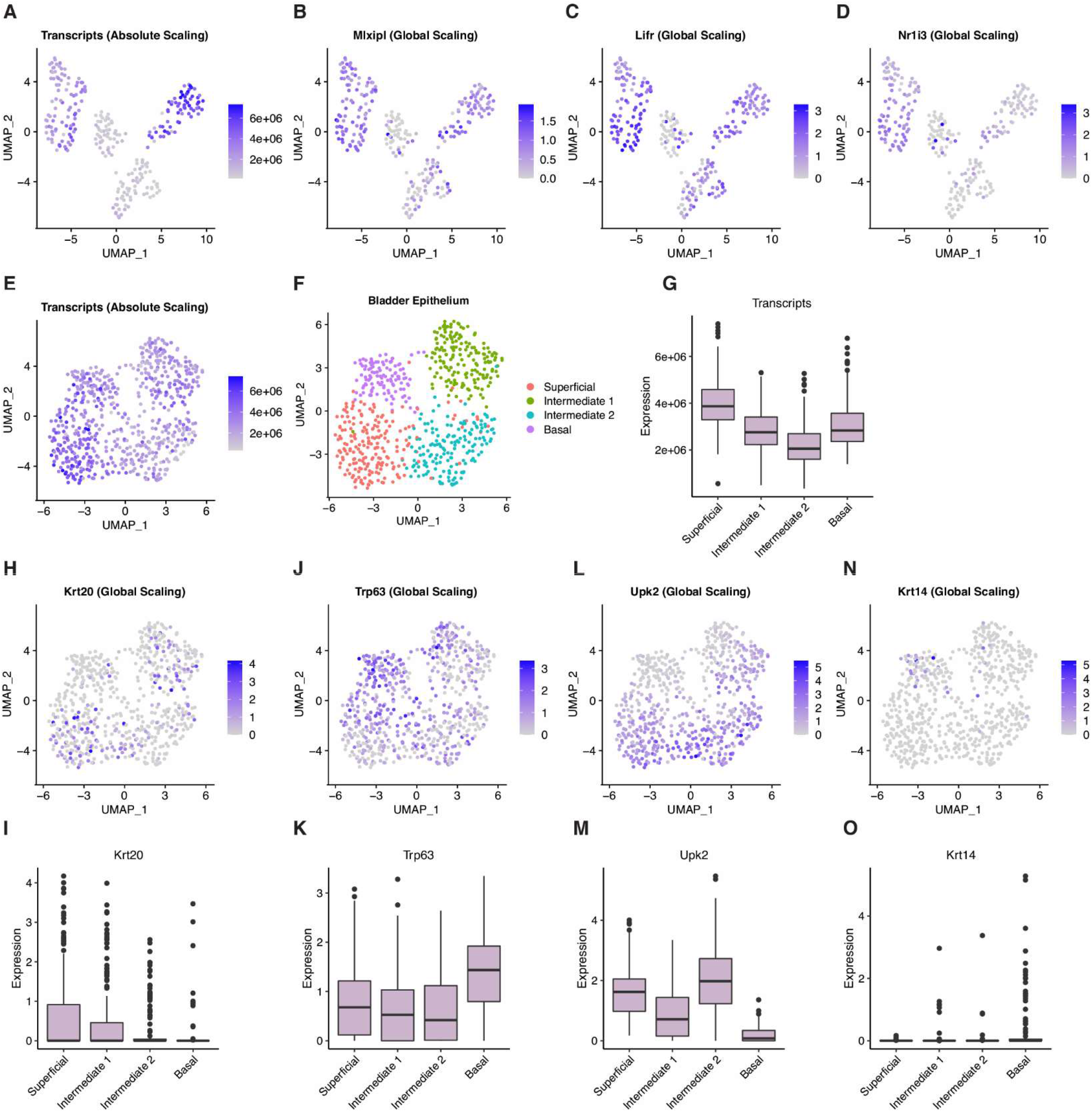
High-Content Cells of the Liver and Bladder Display Polyploidy Markers. **(A-D)** UMAP representation showing ERCC-normalized reads and log2 expression of liver polyploid cell markers. Plots depict dimensionality reduction performed under absolute scaling. **(E-O)** ERCC-normalized transcript abundance and log2 expression of polyploid cell markers in bladder cells. Plots depict dimensionality reduction performed under the indicated scaling type.

**Supplementary Figure 12:**
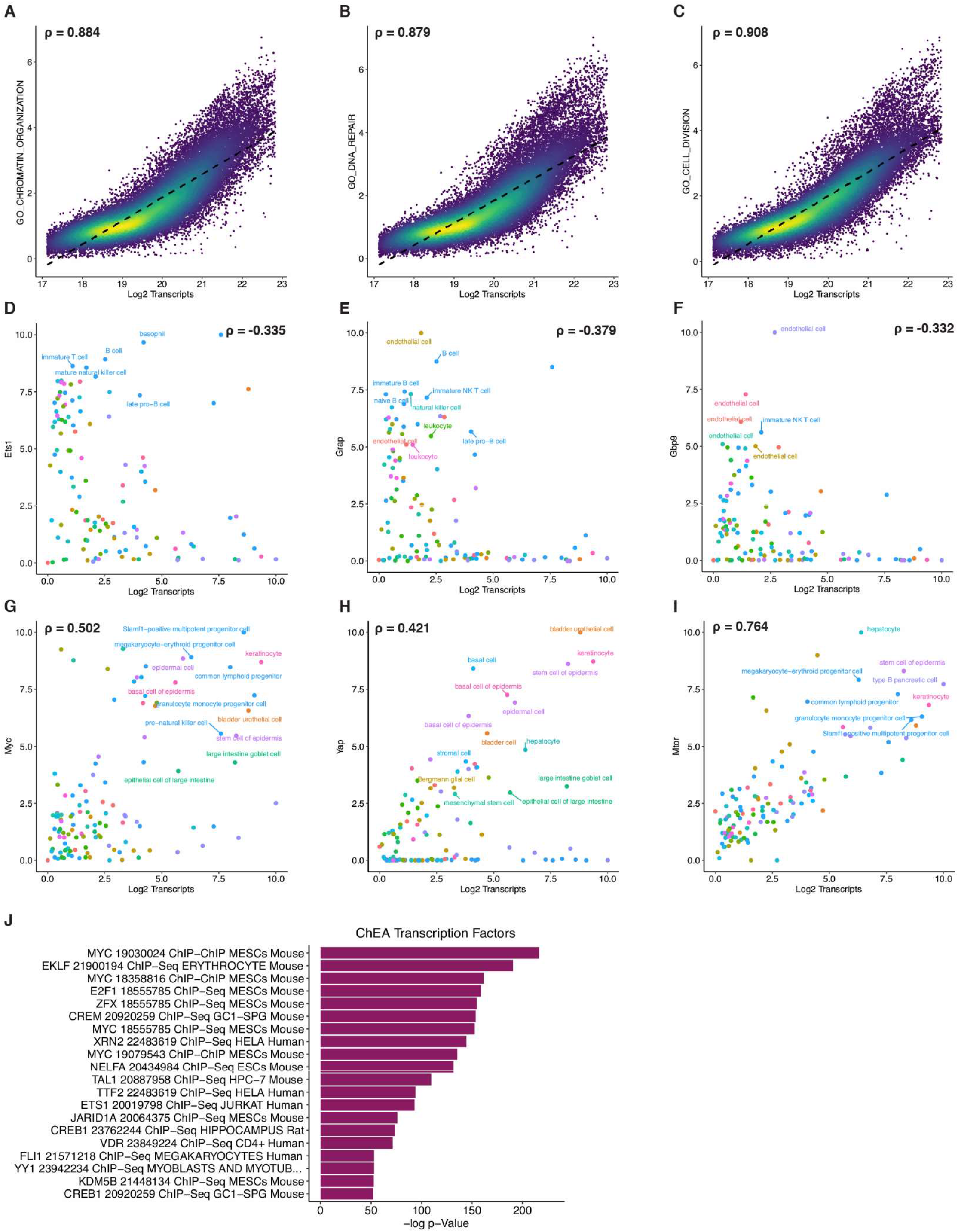
Hypertranscription Hallmarks are Correlated with Transcript Abundance. **(A-C)** Single-cell correlation of transcription signatures with log2 transcript abundance. Signature scores were determined using VISION with absolute-scaled FACS data. Each point represents a single cell, with color scale depicting plotting density. **(D-I)** Positive and negative correlation of select gene expression with log2 transcript abundance. Cells are displayed as averages within cell types, with colors matching organs in (Fig 2). Correlation values represent Spearman coefficients. **(J)** Enrichment analysis for ChEA sets using top 3000 highly correlated genes (to log2 transcript abundance).

**Supplementary Figure 13:**
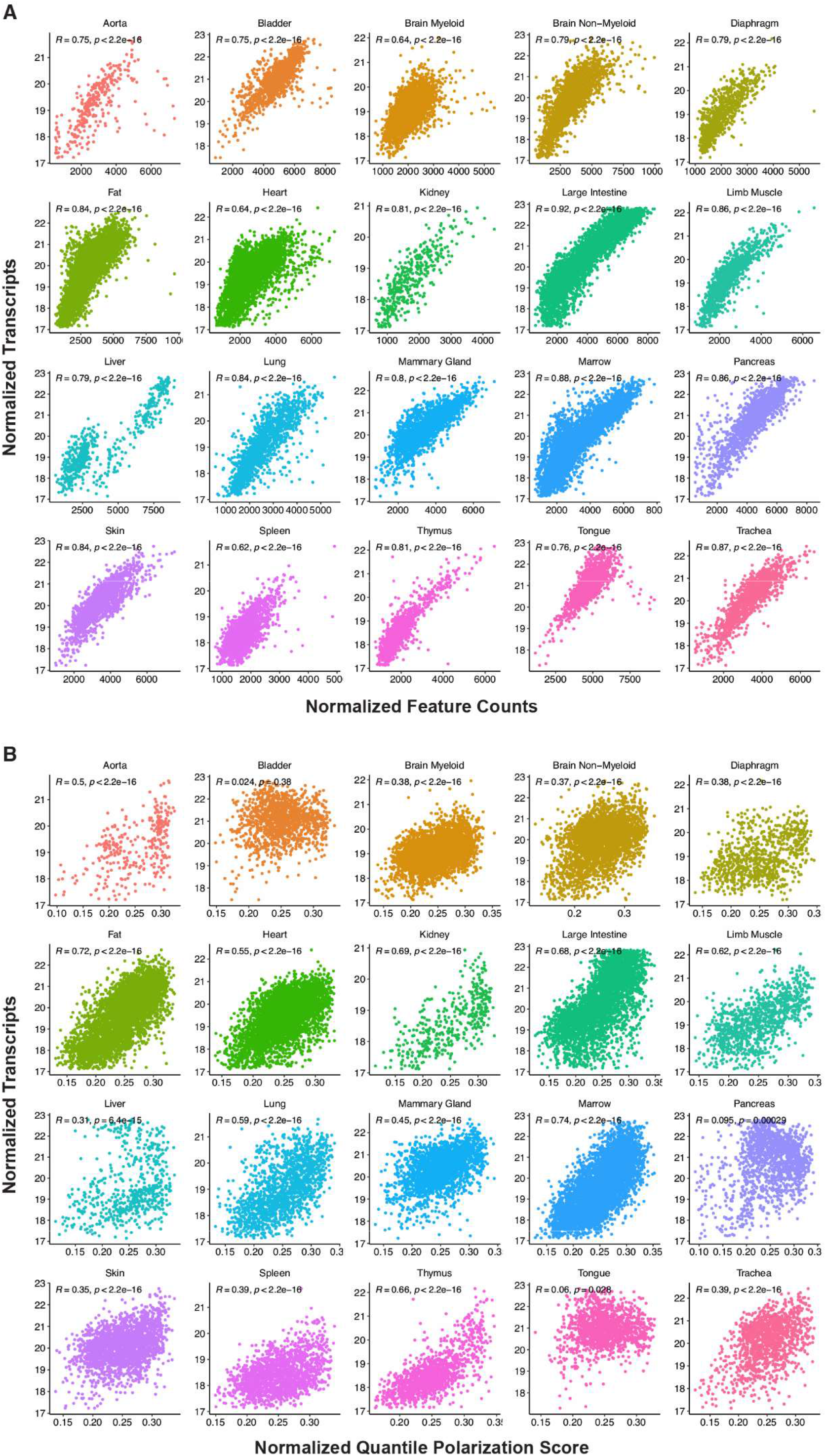
Transcript Content is a Correlated to Markers of Developmental Progression. **(A)** Comparison of log2 absolute-scaled transcript content with feature counts (number of expressed genes). **(B)** Comparison of log2 absolute-scaled transcript content with quantile progression scores, calculated using globally-scaled FACS data. Correlation values in **(A-B)** represent Spearman coefficients.

**Supplementary Figure 14:**
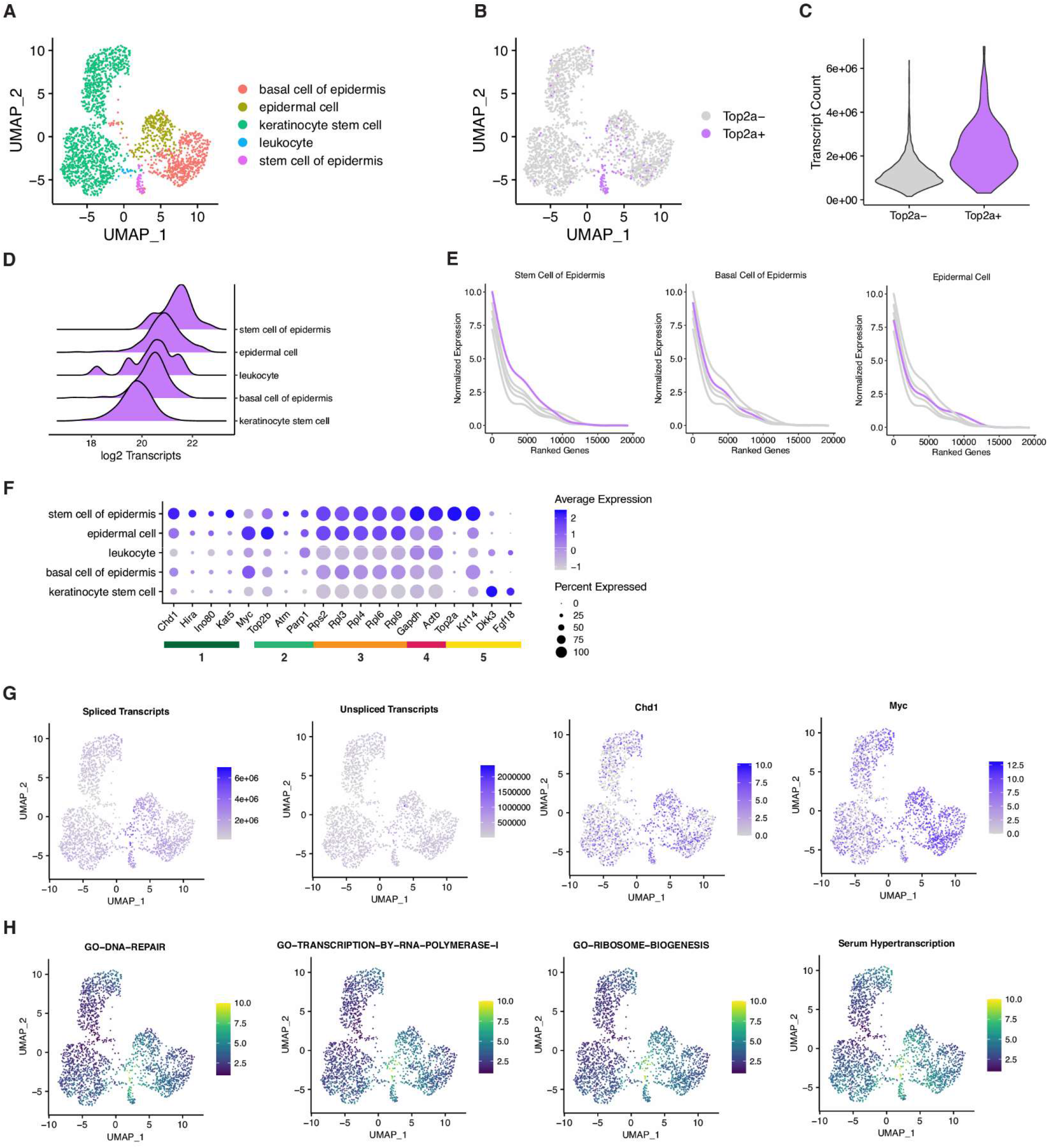
Epidermal Progenitors Display Hallmarks of Hypertranscription. **(A)** UMAP visualization of cell types within the skin FACS dataset. **(B)** Visualization of cycling Top2a+ epidermal stem cells within the dataset. Plots in **(A-B)** depict dimensionality reduction performed under absolute scaling. **(C)** Transcript content differences between Top2a+ and Top2a- cells **(D)** Distribution of cellular transcript content between cell types of the epidermis, ranked by median content. Transcript counts represent log2 ERCC-normalized reads. **(E)** Transcriptome curves depicting gene expression across top 20,000 genes in representative cell types of the epidermis. Individual genes are ranked using combined log2 expression between all cell types. Indicated cell types correspond to the highlighted curve. **(F)** Expression of select genes relevant to hallmarks of hypertranscription, including (1) chromatin remodelers, (2) DNA repair factors, (3) ribosomal genes, (4) housekeeping genes, and (5) epidermal compartment markers. **(G)** UMAP visualization of total spliced/unspliced transcripts, Chd1 and Myc expression under absolute scaling. **(H)** UMAP visualization of signature scores generated by VISION using absolute scaling.

**Supplementary Figure 15:**
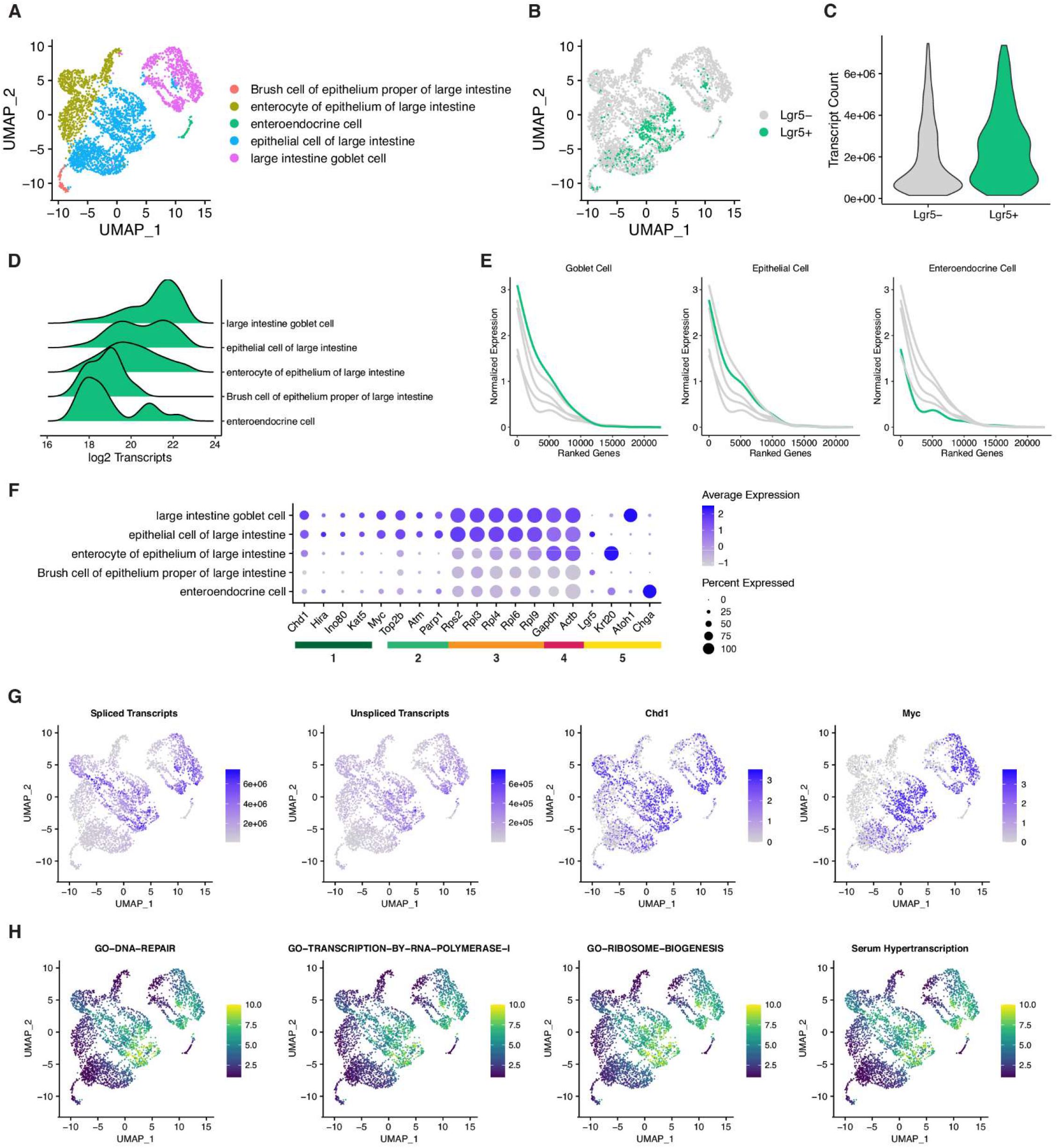
Colonic Epithelium Progenitors Display Hallmarks of Hypertranscription. **(A)** UMAP visualization of cell types within the large intestine FACS dataset. **(B)** Visualization of cycling Lgr5+ intestinal stem cells within the dataset. Plots in **(A-B)**depict dimensionality reduction performed under absolute scaling. **(C)** Transcript content between Lgr5+ and Lgr5- cells. **(D)** Distribution of cellular transcript content between cell types of the colonic epithelium. Transcript counts represent log2 ERCC-normalized reads. **(E)** Transcriptome curves depicting gene expression across top 20,000 genes in representative cell types of the colonic epithelium. Individual genes are ranked using combined log2 expression between all cell types. Indicated cell types correspond to the highlighted curve. **(F)** Expression of select genes relevant to hallmarks of hypertranscription, including (1) chromatin remodelers, (2) DNA repair factors, (3) ribosomal genes, (4) housekeeping genes, and (5) crypt cell markers. **(G)** UMAP visualization of total spliced/unspliced transcripts, Chd1 and Myc expression under absolute scaling. **(H)** UMAP visualization of signature scores generated by VISION using absolute scaling.

**Supplementary Figure 16:**
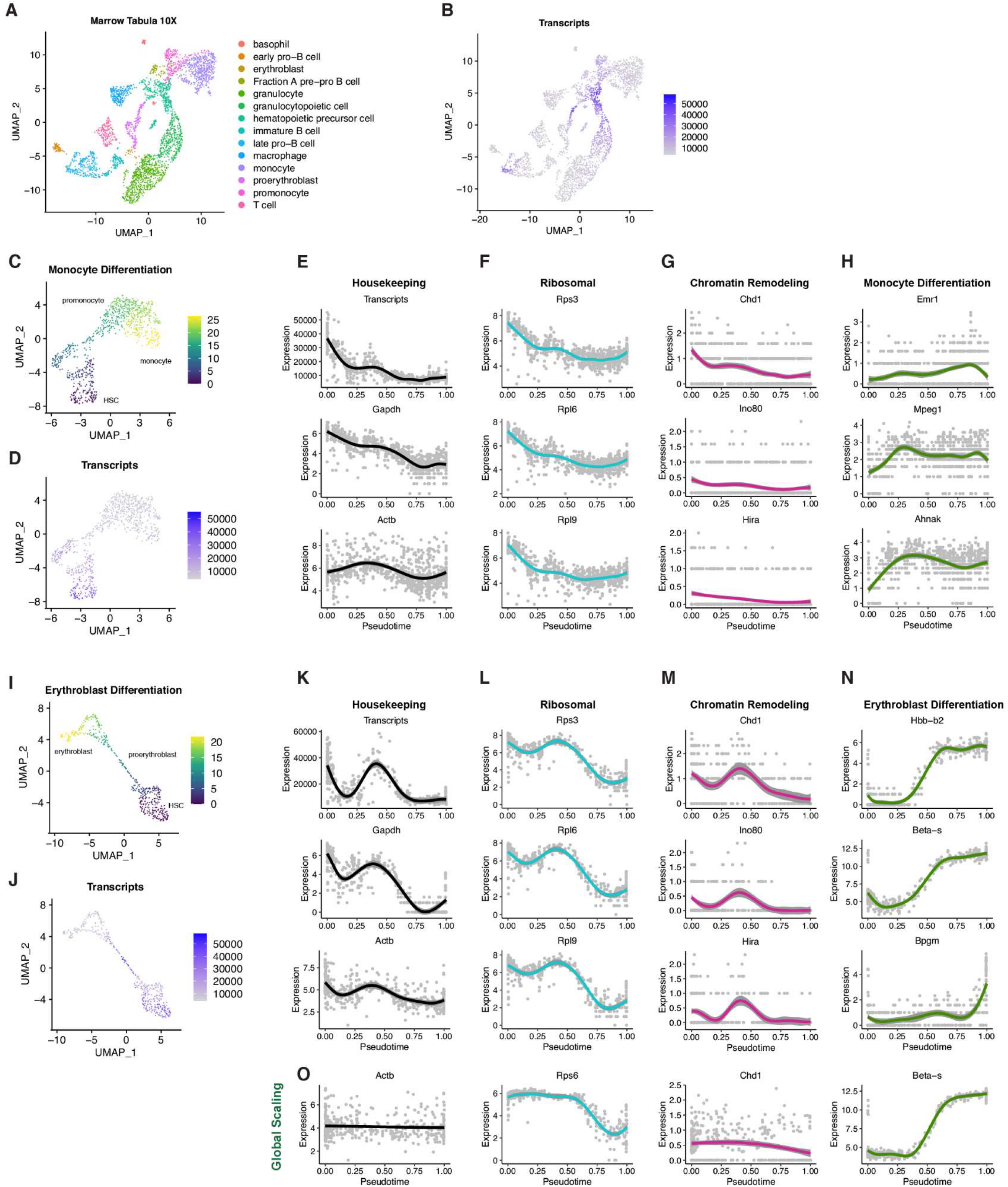
Hypertranscription Marks Developmental Progress Through Erythroblast and Monocyte Differentiation. **(A)** UMAP representation of cell types within Tabula Muris 10X bone marrow dataset. **(B)** Transcript abundance (raw UMIs) across all cells of the bone marrow. **(C-D, I-J)** UMAP visualization of Monocle pseudotime scores and total transcript abundance under absolute scaling. Hematopoietic stem cells were defined as root cells for analysis. Plots in **(A-B), (C-D)**, and **(I-J)** depict dimensionality reduction performed under absolute scaling. **(E-H, K-N)** Gene curves representing absolute-scaled expression of indicated genes through progression of pseudotime. Each point represents single cells ordered by pseudotime values. **(O)** Gene curves representing globally-scaled expression of indicated genes through progression of pseudotime. Each point represents single cells ordered by pseudotime values. Expression values in **(E, K)** represent raw UMIs, values in **(F-H, L-N)** represent log2 raw UMIs.

## SUPPLEMENTARY TABLES

Table S1: Absolute Scaling Statistics for Tabula Muris Atlas

Table S2: Gene-Transcript Content Correlation

Table S3: GO BP and ChEA Terms Enriched in Highly Correlated Genes

